# Development of AI-designed protein binders for detection and targeting of cancer cell surface proteins

**DOI:** 10.1101/2025.05.11.652819

**Authors:** Bianca Broske, Sophie C. Binder, Benjamin A. McEnroe, Tim N. Kempchen, Caroline I. Fandrey, Julia M. Messmer, Elisabeth Tan, Peter Konopka, Dominic Ferber, Michelle C.R. Yong, Marie Kleinert, Alexander Hoch, Katja Blumenstock, Jan M. P. Tödtmann, Johannes Oldenburg, Heiko Rühl, Alexander Semaan, Marieta I. Toma, Kristina Markova, Sebastian Kobold, Tim Rollenske, Matthias Geyer, Stephan Menzel, Tobias Bald, Jonathan L. Schmid-Burgk, Gregor Hagelueken, Michael Hölzel

## Abstract

Artificial intelligence (AI)-based protein design opens new avenues for the rapid generation of new research tools and therapeutics, but experimental validation lags behind the computational design throughput. Here, we present a scalable workflow for the discovery and validation of AI-designed protein binders. Leveraging the RFdiffusion protein design pipeline with a custom filter for stable alpha-helical bundle folds, we construct libraries of thousands of AI-binders against cancer-associated surface proteins. Mammalian cell-surface and phage display screening yield multiple high-affinity PD-L1 binders but fewer hits for CD276 (B7-H3) and VTCN1 (B7-H4), reflecting the target-dependent efficiency of RFdiffusion in generating high-quality designs. Using our experimentally validated AI-designed binder libraries, we benchmark freely available structure prediction models. We find that interface predicted template modelling (ipTM) scores by Chai-1 with ESM embedding correlate well with experimental success and even predict deleterious effects of binding interface mutations. To demonstrate the versatility of AI-binders as research tools, we deploy them in CAR-T cells and also assemble them with fluorophore-labeled streptavidin into tetravalent quattrobinders, which achieve antibody-comparable staining of endogenous PD-L1 by flow cytometry. With high production yields and accessible structural models, AI-designed quattrobinders are versatile and cost-effective research tools amenable to community-driven validation and optimization.

## Introduction

For many decades, structural biology has provided detailed molecular insights into the mechanisms of molecular recognition that govern the interplay between the multitude of large and small molecules within living cells^1^. The corresponding interaction interfaces have been honed by evolution to achieve just the right affinities and specificities. Ever since the first structures of such complexes were determined experimentally, scientists have been thinking about ways to engineer such interfaces artificially, for example by designing or selecting small-molecule inhibitors or antibodies against a target protein.

The advent of artificial intelligence (AI) has dramatically changed structural biology in recent years, and the introduction of AI-based algorithms such as AlphaFold (AF2, AF3)^2,3^ or RoseTTAFold^4,5^ has made it possible to precisely predict protein structures from their amino acid sequence. Additionally, algorithms such as RFdiffusion^6^, AlphaProteo^7^, BindCraft^8^, and Chroma^9^ are able to design either completely artificial proteins or proteins that adhere to user-defined design principles, such as following a particular fold or binding to a target molecule of interest.

A common question is, whether the productive use of these techniques is restricted to specialized labs with long-standing expertise in protein design and access to large-scale computational and screening infrastructure, or, whether other researchers can successfully employ these methods to design proteins that specifically recognize targets that are relevant to their research. Demonstrating that such workflows are accessible and effective beyond expert labs is key to unlocking the broader potential of AI-based protein design.

Here, we have developed an accessible and scalable pipeline to design small artificial protein binders (AI-binders) against three cancer surface targets of the B7-H family that have long been of interest to our labs. For this purpose, we have adapted the open source RFdiffusion and AF2 algorithms to our use case, by implementing custom filter scripts that employ available software tools such as the CCP4^10^ suite or Biopython^11^. Using this pipeline, we designed thousands of trial binders with limited GPU resources and identified functional candidates through a combination of mammalian cell surface display screening, fluorescence-activated cell sorting (FACS) and deep sequencing. The resulting AI-binders were biophysically characterized, and we show how such binders can be rapidly turned into useful research tools in the form of CAR-T cells and open-source detection reagents for flow cytometry.

## Results

### Pipeline for AI-binder design and functional screening

Following the methodology developed by Watson et al.^6^, we generated poly-glycine scaffolds using RFdiffusion, then applied ProteinMPNN (PMPNN)^12,13^ to design amino-acid side chains, and finally “refolded” each sequence with the AlphaFold2 initial-guess algorithm^13^ (**Fig. 1a**). To triage designs, we employed the predicted aligned error (pAE) interaction score from AF2 ^6^, a per-residue-pair confidence metric that estimates the expected positional error for each pair of interface residues. In practice, the mean inter-chain pAE has been shown to be a reliable proxy for interface accuracy, correlating with experimental success^6^.

**Figure 1.**
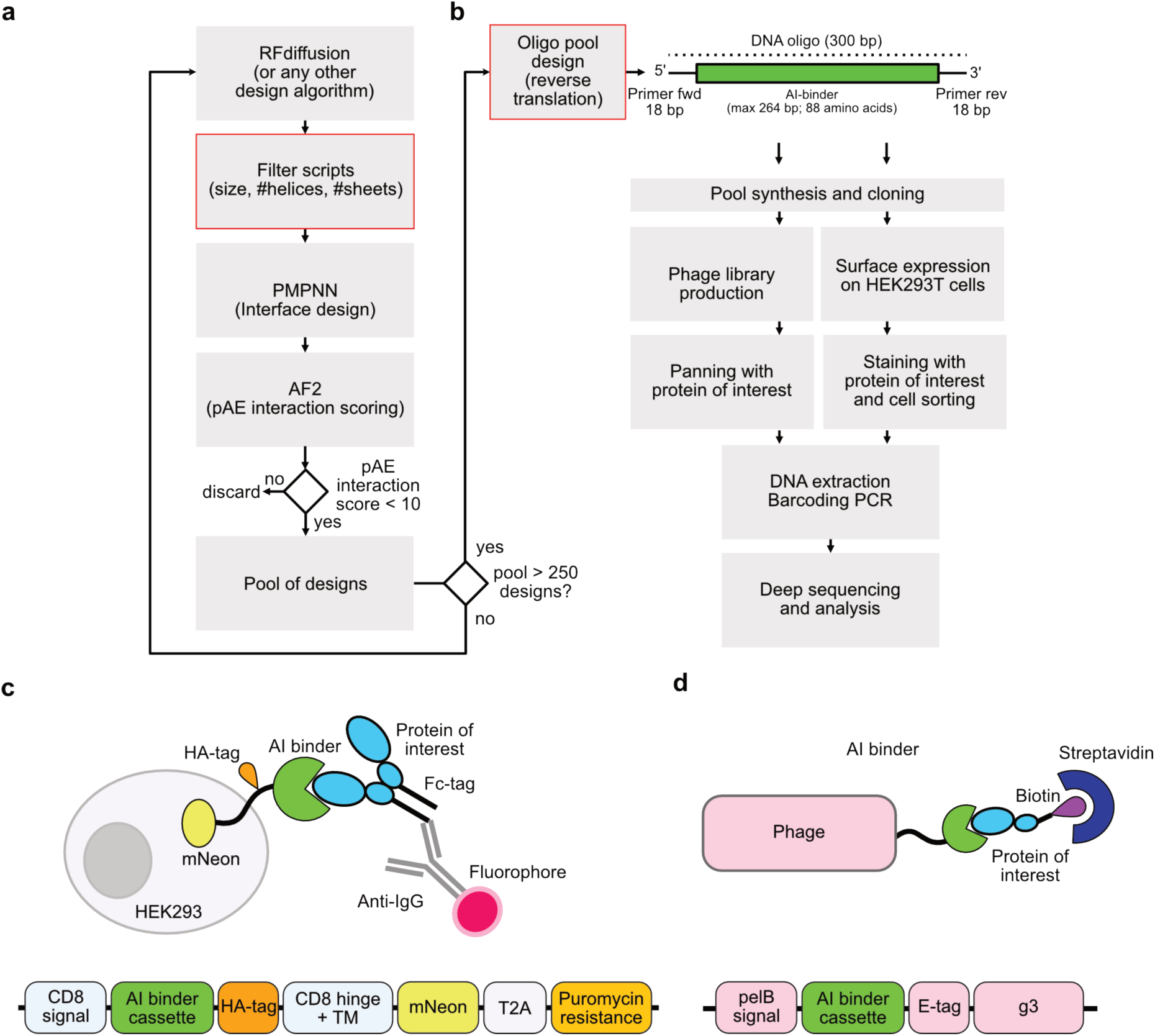
Computational design and functional screening strategy for AI-designed protein binders. **a** Outline of the RFdiffusion design pipeline with structural filtering step. **b** Outline of the functional screening pipelines. **c** Scheme illustrating the principle of mammalian cell-surface display and the construct used for AI-binder presentation. **d** Scheme illustrating the principle of phage display and the construct used for AI-binder presentation.

Depending on the target, RFdiffusion produced designs with good pAE interaction scores (pAE < 10)^6^ with strongly varying efficiency. Due to methodological restrictions (see below), we limited the size of our designs to <= 88 amino acids. We noted that the algorithm often produced alpha-helical hairpins with two rather long alpha helices. Small-scale expression tests indicated poor expression or stability of such designs, probably because these designs do not have a defined hydrophobic core to provide stability. Nevertheless, we noticed that some of the designs had a much more compact fold, similar to affibodies^14^, with three or four short alpha helices that were predicted to form a stable alpha-helical bundles. As we did not want to restrict the freedom of RFdiffusion by forcing it to adhere to a defined fold, we decided to implement a Biopython^11^ script that uses the dssp software^15,16^ to detect secondary structure elements and thereby simply discarded any designs that had less than three alpha-helices. Only the designs that remained after this selection step were fed into the subsequent steps of our pipeline. While this increased the time needed to design a sufficient number of potential AI-binders with promising scores, we were confident that the resulting designs should have a better chance to work in real world applications. Interestingly, we also observed that PMPNN frequently created sequences that exhibited surface-exposed patches of alanines (poly-Ala patches) that lie remotely from the binding interface. It should be noted that our pipeline is not restricted to RFdiffusion but also works with designs that are created with any other algorithm such as Bindcraft^8^ or Chroma^9^.

To identify functional AI-binders at scale, we implemented two pooled screening approaches for targeted enrichment (**Fig. 1b**). For mammalian cell-surface display, we encoded each AI-binder in a chimeric antigen receptor–like construct (**Fig. 1c**), allowing its presentation on the cell surface. Cells bearing functional AI-binders are detected by staining with the target protein and isolated via FACS. In the second approach, phages displaying functional AI-binders are enriched by incubating with the target protein immobilized on streptavidin beads, followed by bead-based purification (**Fig. 1d**). In both workflows, enriched populations are subjected to DNA extraction, and the AI-binder coding sequences serve as templates and identifiers for NGS analysis of PCR-amplified fragments. As mentioned above, we restricted our designed AI-binders to 88 amino acids (264 bp) since 300 bp DNA oligo pools, including both the AI-binder and primer sequences, proved to be the most cost-effective way to generate libraries with thousands of potential candidates for testing. (**Fig. 1b**).

### Design and screening of AI-binders against B7-H family proteins

B7-H family proteins, including PD-L1 (B7-H1), CD276 (B7-H3), and VTCN1 (B7-H4), are well-established surface targets for cancer therapy^17–20^. We set out to design AI-binders against the Ig-like V-type domains leveraging our computational and screening pipeline, using both crystal structures (PD-L1, VTCN1) and AF2 models (CD276) of these surface receptors as input (**Fig. 2a–c**). PD-L1 was included as a benchmarking target, since Watson et al. previously designed AI-binders against PD-L1 using RFdiffusion^6^, allowing us to focus on known hotspot residues. We confirmed that RFdiffusion efficiently generated a large number of candidate PD-L1 binders with promising pAE interaction scores below 10 (about 22 %), and even below 5 in roughly 7 % of the designs (**Fig. 2d**). In contrast, the design of candidate binders against CD276 and VTCN1 was much less efficient, despite testing multiple configurations of hotspot residues, a strategy recommended for increasing yield (Supplementary Table 1). After filtering for designs with at least three alpha-helices, we selected about 250 AI-binder designs for each target (**Fig. 2e**). We also added AI-binder designs from Watson et al.^6^ against PD-L1, TrkA (NTRK1), and insulin receptor (INSR) as controls to obtain a pilot pool with a total of 1,000 designs (Supplementary Tables 2 and 3). HEK293T cells were transduced with the binder library, stained with PD-L1, CD276, or VTCN1 Fc-fusion proteins, and sorted by FACS in biological triplicates (**Fig. 2f**). We detected a 2.5 % PD-L1–positive population, compared with below 1 % for both CD276 and VTCN1. The NGS analyses confirmed these differences, identifying multiple enriched PD-L1 binders, only a handful for CD276, and none for VTCN1. The mNeon positive cell population was used as reference, as all transduced cells expressed mNeon, to normalize for differences in AI-binder representation within the pool. As shown in the scatter plots in **Fig. 2g–i**, enriched AI-binder candidates deviate above the diagonal toward the top of the plot and separate from the main cloud. Notably, our screen also identified the PD-L1 binders previously designed and validated by Watson et al. which were included as positive controls. Phage display generated largely congruent enrichments for PD-L1 binders, and only one additional candidate was identified by phage display that had not been identified by the mammalian cell surface display (**Fig. 2j**). No clear outliers were identified for CD276 and VTCN1 in the phage-display screen, which is consistent with the mammalian cell-surface display results showing only a few candidates for CD276 and none for VTCN1 (Supplementary Fig. 1a, b).

**Figure 2.**
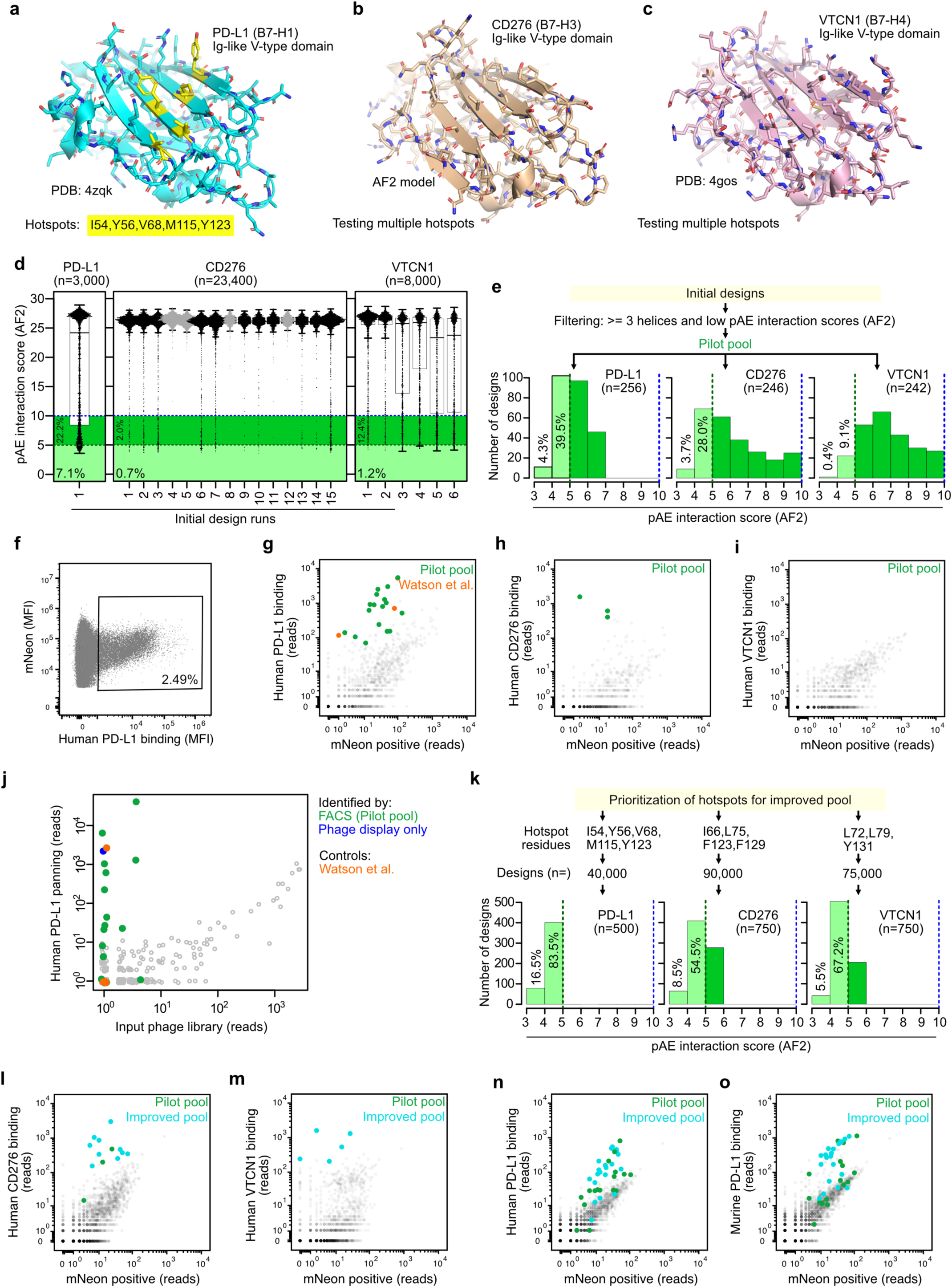
*De novo* design and experimental screening of AI-binders targeting B7-H family proteins. **a** Cartoon representation of the PD-L1 crystal structure (PDB:4zqk), **b** CD276 (B7-H3) AF2 model, and **c** VTCN1 (B7-H4) crystal structure (PDB:4gos). **d** Boxplots with individual data points summarizing AF2 pAE interaction scores of indicated design runs. The total number of designs per target is indicated as well as the percentage of designs with low (good) scores. **e** Histogram depicting the distribution of pAE interaction scores of the best designs used for the pilot pools. **f** Representative scatter plot illustrating a subpopulation of HEK293T cells from the pilot pool binding to human PD-L1 Fc fusion protein. **g** Representative scatter plot showing enriched PD-L1 AI-binders identified by NGS analysis of the human PD-L1-binding subpopulation. **h**, **i** NGS analyses as described in g but results for CD276 and VTCN1. **j** NGS analysis of panning by phage display showing enriched PD-L1 AI-binders. Binders identified by mammalian cell surface display are marked in green, binders only identified by phage display in blue, and positive controls in orange. **k** Histogram depicting the distribution of pAE interaction scores of designs used for the improved pools. **l** Representative scatter plot showing enriched CD276 AI-binders identified by NGS analysis of the human CD276-binding subpopulation from the improved pool. **m** Same analysis as described in l but for VTCN1 AI-binders from the improved pool. **n, o** Same analysis as described in l but for PD-L1 AI-binders from the improved pool binding to human or murine PD-L1.

Given that the CD276 and VTCN1 libraries exhibited higher (i.e., worse) pAE interaction scores than the PD-L1 library, we hypothesized that expanding the number of designs to include more candidates with better scores would improve the success rates for CD276 and VTCN1 (**Fig. 2k**). For the second round of computational designs, we chose the most effective hotspot configuration for CD276 and VTCN1 identified in the first round (Supplementary Fig. 1c, d; Supplementary Tables 4–6). Indeed, screening the improved pool resulted in additional hits for CD276 and a few candidate VTCN1 binders (**Fig. 2l,m**). For PD-L1, we likewise identified additional binders, and comparative staining with a murine PD-L1 Fc fusion protein indicated that our library contains cross-reactive, as well as and human- and mouse-specific PD-L1 binders, respectively (**Fig. 2n,o**).

### Characterization of de novo designed PD-L1 binders

**Fig. 3a** shows the predicted structures of representative PD-L1 binders from the pilot pool in complex with the Ig-like V-type domain of PD-L1. For validation, individual PD-L1 binders were expressed on HEK293T cells, including the positive controls from Watson et al. (WS_76, WS_8, WS_5, WS_45)^6^, and assessed by flow cytometry for binding to human and murine PD-L1 (**Fig. 3b**; Supplementary Fig. 2a). Efficient cell surface expression of AI-binders was confirmed by detection of the HA-tag incorporated in the extracellular part of the construct (Supplementary Fig. 2b; **Fig. 1c**). Underscoring the robustness of our screening platform, 15 of 17 PD-L1 binders confirmed binding to human PD-L1. Interestingly, several were cross-reactive with the murine ortholog of PD-L1.

**Figure 3.**
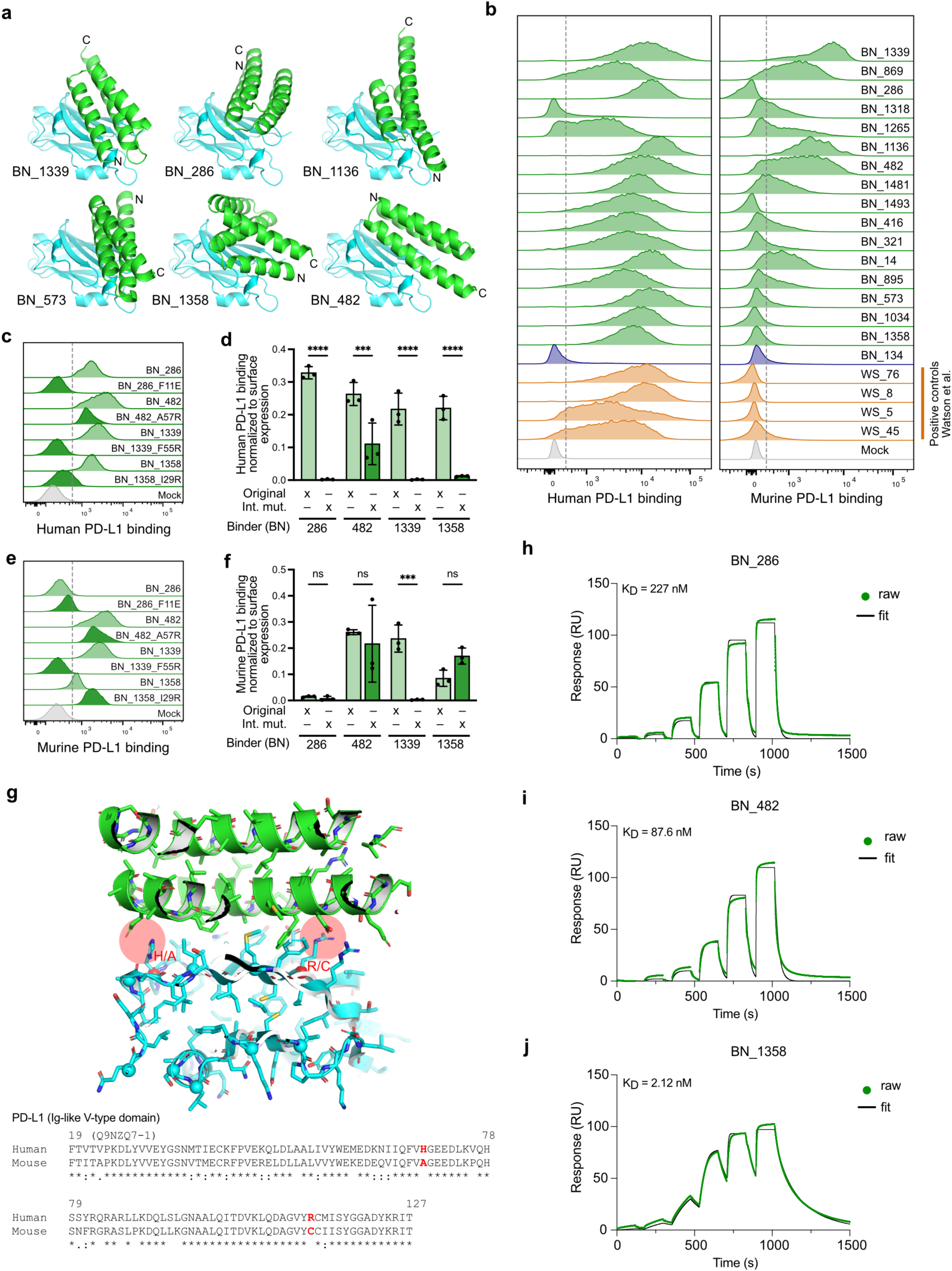
Characterization of *de novo* designed PD-L1 AI-binders reveals both species selectivity and cross-reactivity. **a** Cartoon representation of the predicted complexes between PD-L1 AI-binders and the Ig-like V-type domain of human PD-L1. N, C: N- or C-termini. **b** Flow cytometric analyses evaluating the binding of individual PD-L1 binders to human and murine PD-L1 Fc-fusion proteins by HEK293T cell surface display. Binders identified by mammalian cell surface display are marked in green, binders only identified by phage display in blue, and positive controls in orange. **c** Experiment and analysis as described in b but for PD-L1 AI-binder interface mutants binding to human PD-L1. **d** Quantification of the experiment in c from biological triplicates (n=3). Mean fluorescence intensity (MFI) is normalized to HA-tag signal (cell surface expression of AI-binder). **e** Experiment and analysis as described in c but for PD-L1 interface mutants binding to murine PD-L1. **f** Quantification of the experiment in e from biological triplicates (n=3). **g** Cartoon representation with side chains of the predicted complex between PD-L1 binder BN_286 (green) and human PD-L1 (cyan). Residues highlighted in red might explain selective binding to human but not murine PD-L1. **h** SPR (Surface Plasmon Resonance) sensorgrams showing raw binding data (green) and fitted curves (black) used to determine binding affinities (KD) of PD-L1 binder BN_286 binding to human PD-L1. **i** Same analysis as described before but for BN_482, and **j** for BN_1358. ***p<0.001; ****p<0.0001; two-way ANOVA with Šídák’s multiple comparisons test.

To confirm that the designed interfaces formed as intended, we introduced non-conservative interface mutations into four of our PD-L1 binders and tested whether these alterations could disrupt their interaction with their target. The resulting AI-binder interface mutants were again expressed on the surface of HEK293T cells, stained with the human PD-L1 Fc-fusion protein and analyzed by flow cytometry (**Fig. 3c,d**). Interface mutations abolished human PD-L1 binding in three of the four AI-binders (BN_286_F11E, BN_1339_F55R, BN_1358_I29R), while the fourth showed only a partial reduction in binding (BN_482_A57R). We also stained with the murine PD-L1 Fc-fusion protein and obtained similar results for BN_1339_F55R and BN_482_A57R but interestingly observed improved binding to murine PD-L1 for BN_1358_I29R (**Fig. 3e,f**). In addition, we also confirmed selective binding of BN_286 to human PD-L1, which is supported by inspection of the predicted structure of the complex, revealing that two prominent interactions are not possible with murine PD-L1 (**Fig. 3g**).

In order to thoroughly study the biophysical properties of our AI-binders, we expressed the proteins in *E. coli*. We were worried that the beforementioned surface alanine patches might lead to aggregation of the proteins during expression and mutated some of the alanine sidechains to polar residues. Supplementary Fig. 3a shows that the AI-binder proteins, including the original (“wild-type”) variants with the alanine patches, expressed very well in *E. coli*. Analytical gel filtration experiments during the purification yielded peaks at the expected elution volumes for the monomeric proteins with no visible aggregates (Supplementary Fig. 3b). We conducted nanoscale differential scanning fluorimetry (nanoDSF) experiments of Trp or Tyr containing AI-binders BN_482 and BN_1358 to determine their thermal stability at elevated temperatures and found that the designs were very stable and even refolded when the temperature was reduced again (Supplementary Fig. 3c). To confirm this observation and to further assess the thermal stability of the designed proteins, we heated lysates of *E. coli* that overexpressed the AI-binders at 94 °C for 20 minutes and found substantial amounts of AI-binders in the cleared supernatant of all constructs (Supplementary Fig. 3d). Next, we used surface plasmon resonance (SPR) spectroscopy to determine the binding characteristics. We immobilized human PD-L1 to the sensor surface and injected concentration series of the binders BN_286, BN_482, and BN_1358 purified from *E. coli*. As expected from flow cytometry, we detected strong interactions between PD-L1 and the AI-binders with dissociation constants in the range of 2 nM to 200 nM (**Fig. 3h–j**). In this assay, BN_1358 was the best binder (K_D_ = 2 nM) with a comparably slow off-rate, compared to binders BN_482 (K_D_ = 87 nM) and BN_286 (K_D_ = 227 nM). Notably, replacing the poly-Ala patches outside the binding interface with hydrophilic residues resulted in modest changes in both the association and dissociation rates compared to the original AI-binders (Supplementary Fig. 4a–c).

### PD-L1 AI-binder validation using CAR-T cells

Next, we set out to leverage the chimeric antigen receptor (CAR)-T cell approach for evaluation of our RFdiffusion *de novo* designed PD-L1 AI-binders. CAR-T cells recognize surface antigens on cancer cells and therefore provide a complimentary approach for evaluating AI-binder functionality as recently shown by others^21^. The PD-L1 AI-binders BN_1358, BN_482, and BN_286 were cloned into a second-generation CAR expression construct with an additional HA-tag for detection of surface expression, and the RQR8 peptide separated by a P2A cleavage site (Supplementary Fig. 5a). RQR8 is displayed on the cell membrane and recognized by a CD34-specific antibody (QBEND10)^22^, enabling straightforward monitoring and enrichment of T cells transduced with the CAR construct (Supplementary Fig. 5b). Upon co-culture, T cells expressing the PD-L1 binder CAR selectively recognized CHO (Chinese hamster ovarian) target cells overexpressing human PD-L1 compared to controls determined by intracellular cytokine staining (Supplementary Fig. 5c,d). Importantly, PD-L1 binder CAR T cells also selectively recognized endogenously expressed PD-L1 on different cancer cells lines compared to matched CRISPR-Cas9 engineered PD-L1 knockout (KO) cells (Supplementary Fig. 5e,f). Together, these findings demonstrate that RFdiffusion-designed AI-binder CAR-T cells can effectively target endogenously expressed cancer surface proteins.

### CAR-T cells with CD276 AI-binders

Encouraged by these results, we applied the CAR-T cell platform to assess our AI-binders against CD276 (B7-H3). The extracellular part of CD276 consists of an Ig-like V-type (V) and Ig-like C2-type (C) domain. Because humans, but not rodents, harbor an exon duplication, human cells express a longer isoform of CD276 (V–C–V–C, also known as 4Ig CD276), although some cells also produce the shorter, two-domain (V–C) variant (2Ig CD276)^23^ (**Fig. 4a**). First, we validated the functionality of our CD276 AI-binders by cell surface display, assessing their binding to both the 2Ig and 4Ig CD276 isoforms produced as Fc-fusion proteins (**Fig. 4b,c**). Interface mutants again confirmed that the designed interfaces formed as intended, abolishing binding to 2Ig and 4Ig CD276 in all cases. From this experiment, we selected the three most promising CD276 AI-binder candidates (BN_616-606, BN_486-1519, and BN_616-2162) for further evaluation by our CAR platform. A recent study also reported AI-designed CD276 binders (MiB, HM9), though a different design algorithm was used^24^, followed by multiple rounds of experimental affinity maturation^21^. Of note, a picomolar binding affinity was reported for HM9, which is why we included the MiB and HM9 CD276 AI-binders as positive controls in our assay. Transduction of T cells was successful for all five CAR constructs achieving comparable efficiencies. However, we noted markedly reduced surface expression of MiB and HM9 containing CARs compared to our CD276 AI-binder CARs (**Fig. 4d–f**). All five CD276 AI-binder CAR-T cells selectively recognized CHO cells overexpressing either 2Ig or 4Ig CD276, though HM9 CAR-T cells were also activated by control CHO cells (Supplementary Fig. 6a–c). Since HM9 was the only AI-binder that cross-reacted with murine CD276 (Supplementary Fig. 6d), we believe that HM9 CAR T-cell recognition of CHO cells is a cross-species target-specific binding rather than an off-target effect. This conclusion is further supported by the structural models showing that all sequence differences between murine and hamster (*C. griseus*) CD276 lie outside the predicted binding interface (Supplementary Fig. 6e).

**Figure 4.**
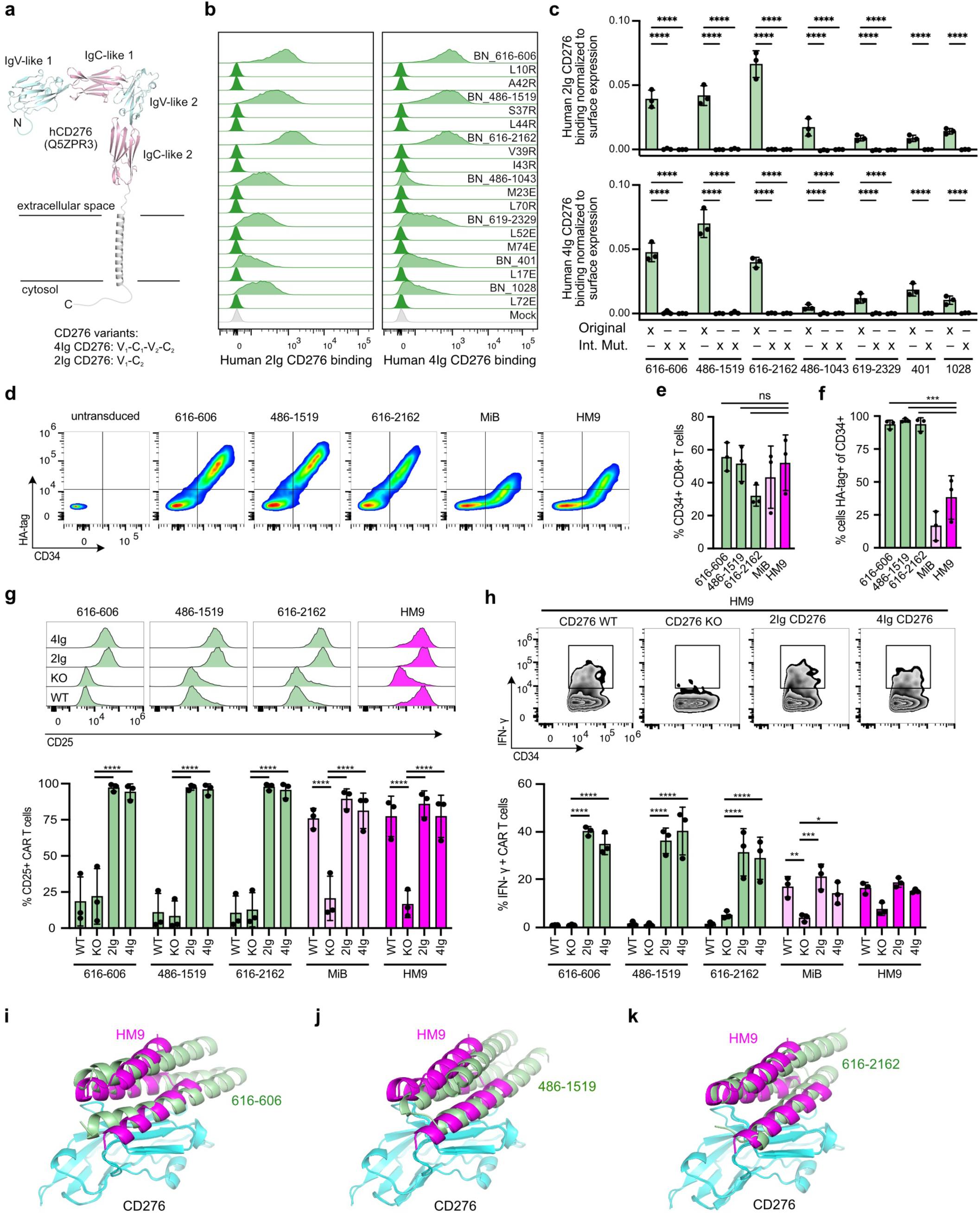
CD276 AI-binders deployed in CAR-T cells reveals critical roles of binding affinity and surface expression for endogenous target recognition. **a** AlphaFold model of CD276 and domain architecture of human 2Ig and 4Ig CD276 isoforms. V: Ig-like V-type domain; C: Ig-like C2-type domain. **b** Flow cytometric analyses evaluating the binding of individual CD276 AI-binders, and corresponding interface mutants, to human 2Ig and 4Ig CD276 Fc-fusion proteins by HEK293T cell surface display. **c** Quantification of the experiment described in b performed in biological triplicates (n=3). **d** Representative flow cytometric plots showing surface expression of RQR8 and CARs (HA-tag) of indicated CD276 AI-binders. **e** Corresponding quantification of transduction efficiency from experiments performed in biological triplicates (n=3). **f** Corresponding quantification of AI-binder CAR surface expression on transduced RQR8+ T cells. Experiments performed in biological triplicates (n=3). **g** Flow cytometric analyses of CD25 surface expression on indicated CD276 AI-binder CAR-T cells co-cultured with SK-MEL-28 CD276 WT, KO and 2Ig, 4Ig CD276 isoforms reconstituted SK-MEL-28 CD276 KO cells, respectively, with corresponding quantification (n=3). **h** Experiments as described in g. Representative flow cytometric plots show IFN-γ induction in HM9 CD276 CAR-T cells as an example and corresponding quantification of indicated CD276 AI-binder CAR-T cells (n=3). **i** Cartoon representation of AF3 structure models showing overlay of HM9 and BN_616-606 CD276 AI-binders bound to the Ig-like V-type domain of human CD276. **j** As described in i but for BN_486-1519, and **k** BN_616-2162. *p<0.05; **p<0.01; ***p<0.001; ****p<0.0001; ns, non-significant; two-way ANOVA with Šídák’s multiple comparisons test.

We then co-cultured the various CD276 AI-binder CAR-T cells with human SK-MEL-28 melanoma cells, which endogenously express CD276. In addition, we used CRISPR-Cas9 to generate matched SK-MEL-28 CD276 knockout (KO) cells and re-expressed either 2Ig or 4Ig CD276 at levels higher than endogenous CD276 expression in SK-MEL-28 wild-type (WT) cells (Supplementary Fig. 6f). All five CD276 AI-binder CAR-T cells were selectively activated by 2Ig or 4Ig CD276 overexpressed on SK-MEL-28 CD276 KO cells (**Fig. 4g,h**), though MiB and HM9 CAR-T cells produced less IFN-γ (**Fig. 4h**). Notably, only MiB and HM9 CAR-T cells responded to endogenously expressed CD276 on SK-MEL-28 WT cells by CD25 upregulation and IFN-γ production, presumably due to high binding affinity of MiB and HM9 as previously reported. Overlaying the structural models shows that our CD276 AI-binders and HM9 occupy largely the same binding site on CD276 (**Fig. 4i-k**). From these findings, we conclude that deploying AI-binders in CAR-T cell platforms requires careful evaluation of multiple factors, in particular target binding affinity and efficient cell-surface expression. In this context, incorporating mammalian cell-surface display during binder discovery helps to select AI-binders with the biochemical profiles needed for effective expression on CAR-T cells.

### Computational scores versus experimental success rates

Using RFdiffusion, the efficiency and success rate of designing functional AI-binders varied widely among our three targets, although they all belong to the same family of B7-H proteins related (about 30 % amino acid identity in their Ig-like V-type domains). Thus, we asked how well different in silico interaction scores predict experimental binding and if they could help to streamline the pipeline. Besides the AF2 pAE interaction score generated during the initial design with RFdiffusion, we rescored the designs with two recently released open-source models, Chai-1^25^ and Boltz-1^26^. **Fig. 5a** outlines our computational re-scoring strategy that uses protein sequences (FASTA format) as input data, which are also extracted from PDB files. Multiple sequence alignments (MSA) for Boltz-1 were locally generated with Mmseq2 in GPU mode^27^. We used Chai-1 primarily with the ESM large language model (LLM) embedding^28^. Both models generate ipTM scores (interface predicted template modelling) that range between 0 (worst) and 1 (best), where ipTM scores higher than 0.7 are considered promising. All designs from the PD-L1, CD276, and VTCN1 pools were ranked based on the respective scores from best to worst (**Fig. 5b**). Highlighting the experimentally validated AI-binders indeed showed a clear correlation between better scores and experimental success, though this analysis was limited for VTCN1 due to the low numbers of functional AI-binders (Supplementary Fig. 7). Of note, the MiB and HM9 CD276 binders were ranked highest by all three models. In a side-by-side comparison, the Chai-1 ESM ipTM score showed the highest normalized enrichment scores (NES), and thus best performance, albeit the difference was modest (**Fig. 5c,d**).

**Figure 5.**
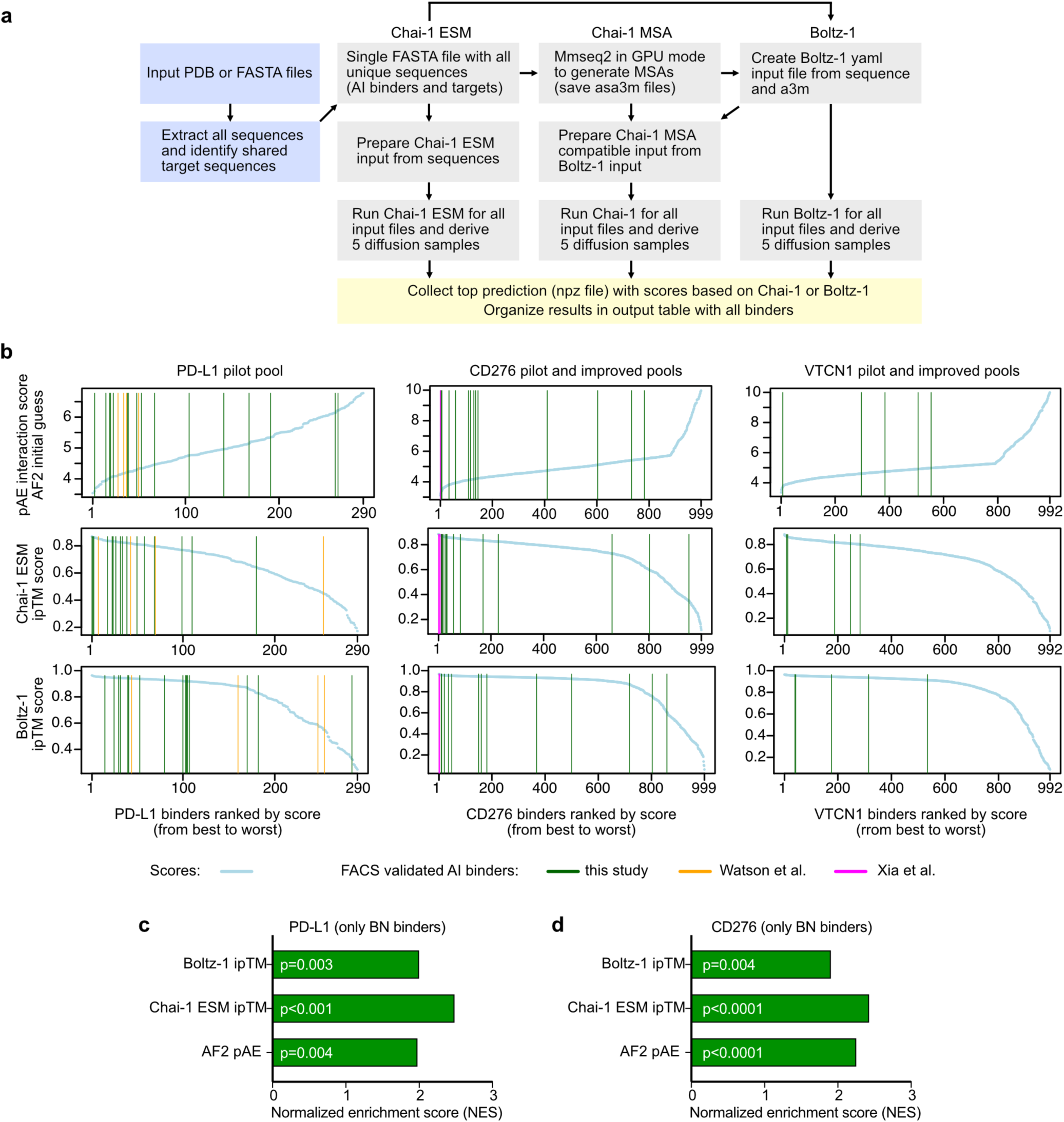
Association between interaction scores generated by different open-source structure prediction models and experimental success of AI-binders. **a** Outline of computational strategy for rescoring of AI-binder pools and generation of iptm scores by Chai-1 and Boltz-1. **b** Enrichment plots visualizing correlation between depicted interaction scores and experimental success. AI-Binders from indicated pools ranked from left to right by interaction scores (best to worst). Colored vertical lines indicate functional AI-binders from our study (green), or from other studies (orange, magenta). **c** Normalized enrichment scores (NES) reflecting the association between interaction scores and experimental success in the PD-L1 AI-binder pilot pool. **d** Same analysis as in c but for the CD276 AI-binder pool (combined pilot and improved pool).

### Prediction of interface mutation effects by Chai-1

Building on our rescoring results, we examined in depth the concordance between the AF2 pAE interaction- and the Chai-1 ESM ipTM scores. Albeit both scores correlate well in general, we also noted some interesting cases of substantial disagreement when analyzing the scores of the PD-L1 pilot pool binders. As highlighted in **Fig. 6a**, this discrepancy was of particular interest for our experimentally validated PD-L1 binders, which roughly segregated into two discordant groups, A and C, and a concordant group B. Integration of normalized flow cytometry data revealed that group C binders exhibited higher binding signals than group A, suggesting that the Chai-1 ESM ipTM score outperforms the pAE interaction score (as implemented in the RFdiffusion design pipeline) as a predictor of experimental binding strength, at least in our limited dataset (**Fig. 6b**). In this context, BN_1358 was a notable example, given its high affinity (K_D_ = 2 nM) determined by SPR analysis (**Fig. 3j**), the relatively high AF2 pAE interaction score of 6.2, and the good Chai-1 ESM ipTM score of 0.84. Overlay of the complexes predicted by AF2 initial guess within RFdiffusion and Chai-1 ESM showed a marked shift of the first alpha helix, with Chai-1 ESM predicting a more compact conformation of BN_1358 (**Fig. 6c**).

**Figure 6.**
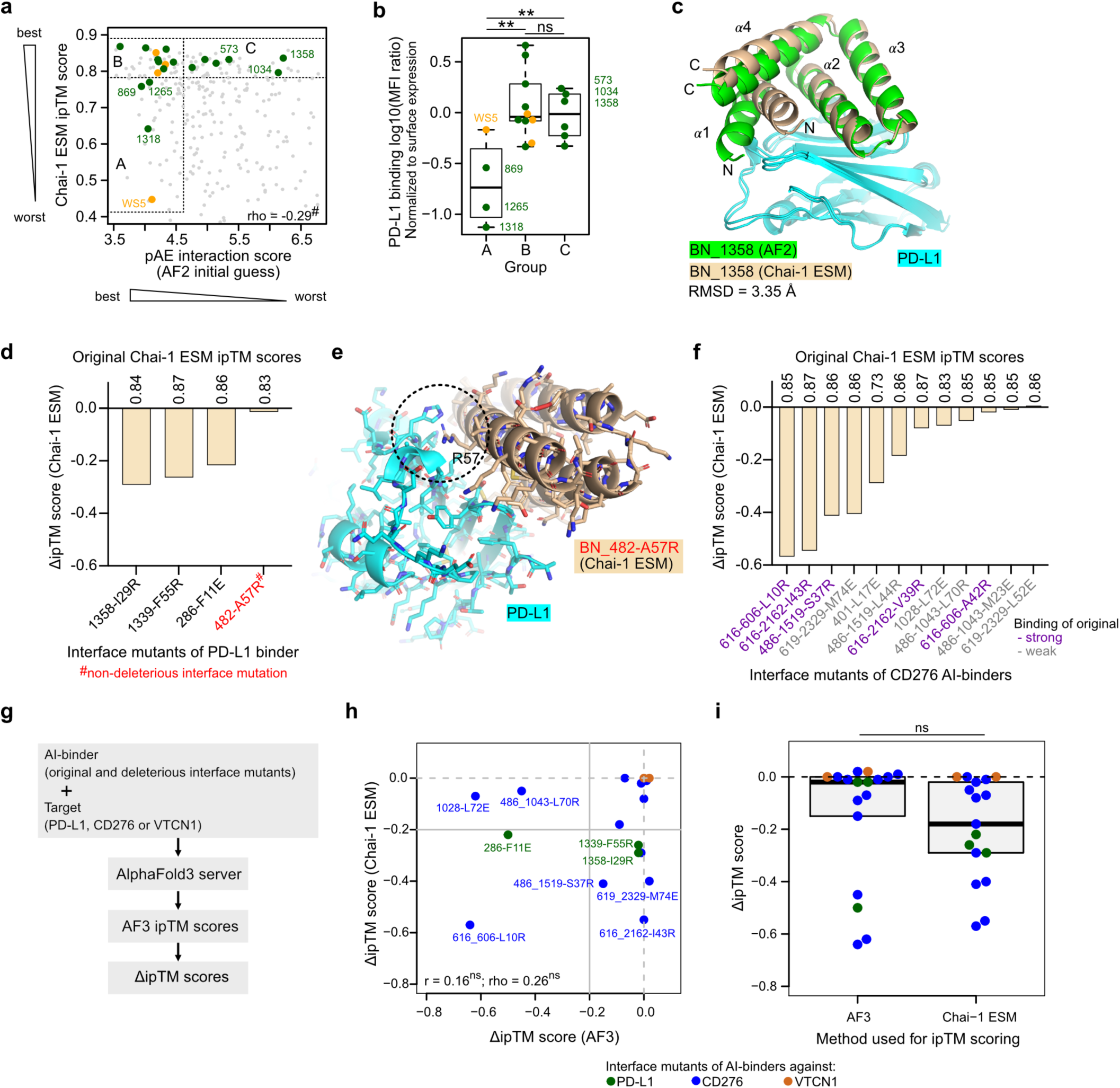
Chai-1 interaction scores predict deleterious effects of binding interface mutations. **a** Scatter plot showing the correlation between pAE (AF2) and ipTM (Chai-1 ESM) interaction scores for the PD-L1 pilot pool. Functional PD-L1 AI-binders from our study highlighted in green, and some labelled with identifier. Positive controls from other study in orange. Rho: Spearman’s rank correlation coefficient. ^#^p=6.38e-07. Groups A, B, and C indicate concordant or discordant scores. **b** Log10-transformed mean fluorescence intensity (MFI) ratio. PD-L1 binding normalized to AI-binder surface expression (Ha-tag) by different groups. **FDR<0.01, two-sided pairwise t-test. **c** Overlay of BN_1358 bound to PD-L1, shown in cartoon representation, with structures predicted by AF2 (green) and Chai-1 with ESM embeddings (beige). Root mean square deviation (RMSD) determined for binder structure predictions only. Å, Ångström. α-helices are numbered from N-terminus onwards. N, C: N- or C-termini. **d** Change in Chai-1 ESM ipTM score caused by interface mutations in PD-L1 AI-binders. **e** Cartoon representation of the BN_A482-A57R interface mutant in complex with Ig-like V-type domain of human PD-L1, as predicted by Chai-1 with ESM embeddings. The outward position of R57 is highlighted with a dashed circle. **f** Change in Chai-1 ESM ipTM score caused by interface mutations in CD276 AI-binders. **g** Outline of strategy to determine AlphaFold3 (AF3) ipTM scores and corresponding ΔipTM scores. **h** Scatter plot showing relationship between AF3 and Chai-1 ESM ΔipTM scores. r, Pearson’s correlation value; rho, Spearman’s rank correlation value; ns, not significant. **i** Boxplot comparing AF3 and Chai-1 ESM ΔipTM scores. Two-sided Wilcoxon-test; ns, not significant.

Intriguingly, rescoring of PD-L1 binder interface mutants using Chai-1 ESM revealed that three of the four mutants exhibited a clear decrease in their ipTM scores (ΔipTM) relative to their matched original PD-L1 binders (**Fig. 6d**; Supplementary Table 7). Notably, the fourth mutant (BN_482_A57R) retained an ipTM score comparable to the “wild-type”, original binder and, in line with this prediction, had shown significant residual binding activity by flow cytometry (**Fig. 3c,d**). According to the Chai-1 prediction, R57 undergoes an outward twist instead of imposing the intended steric hindrance at the binding interface, giving a possible explanation for our experimental observation **(Fig. 6e**). When rescoring the CD276 interface mutants, we observed reduction in ipTM scores for 11 of the 12 mutants, with changes (ΔipTM) ranging from 0.01 (BN_486-1043_M23E) to 0.57 (BN_616-606_L10R), possibly reflecting differences in the binding strength of the matched original AI-binders and accuracy of the initial prediction (**Fig. 6f**).

We then compared ΔipTM scores for deleterious interface mutants predicted by AlphaFold3 and Chai-1 ESM (**Figure 6g**). Interestingly, the two sets of ΔipTM scores did not significantly correlate, highlighting differences in the prediction methodologies (**Figure 6h**). Nonetheless, both AF3 and Chai-1 ESM ΔipTM scores captured the deleterious impact of interface mutations, though to a variable degree, with Chai-1 ESM ΔipTM scores being slightly more pronounced (**Figure 6i**).

### Quattrobinders as flow cytometry reagents

The possibility to design specific, high affinity AI-binders has many practical implications and may help addressing long-standing problems, such as the widespread demand across the research community for reliable open-source protein detection tools^29–31^. These are particularly valuable for applications like flow cytometry experiments. Hence, we decided to assess the usability of AI-binders as staining reagents. To enable site-specific biotinylation, we introduced a single cysteine residue into the AI-binder surface, opposite of its binding interface. The modified AI-binders were then expressed in *E. coli* (Supplementary Fig. 3a), biotinylated, and complexed with fluorophore-conjugated streptavidin to form a tetravalent staining reagent, which we have termed “quattrobinder” (**Fig. 7a**). Expression of the majority of our cysteine-engineered AI-binders in *E. coli* proved highly efficient, producing up to 120 mg per 2 L of culture. We selected four of our best PD-L1 binders for quattrobinder formation, as well as one of our poorest PD-L1 binders (BN_1265) to span a broad performance range. Then, we stained CHO cells overexpressing human PD-L1 with either AF647-labelled PD-L1 quattrobinders or an AF647-labelled PD-L1 antibody. As shown in **Fig. 7b**, our best PD-L1 quattrobinders produced signal intensities on par with the commercial antibody and exhibited negligible background by flow cytometry.

**Figure 7.**
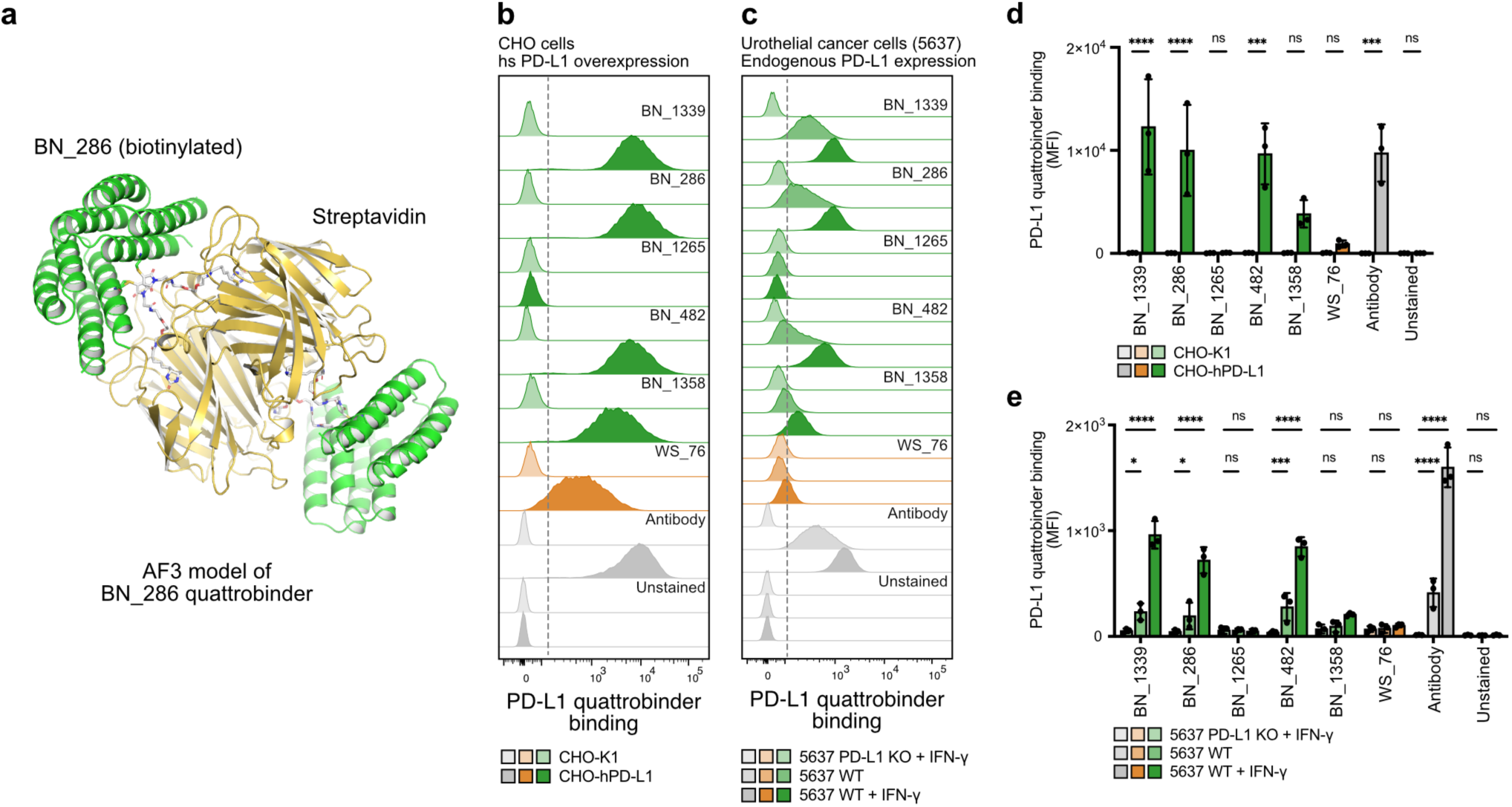
PD-L1 AI-binders assembled as fluorescent quattrobinders specifically detect endogenous PD-L1 expression on cancer cells. **a** Cartoon representation of the predicted structure of the biotinylated PD-L1 AI-binder BN_286 assembled as quattrobinder. **b** Flow cytometric analyses showing detection of human PD-L1 overexpressed on CHO cells by AF647-conjugated PD-L1 quattrobinders compared to an AF647-conjugated anti-PD-L1 antibody. **c** Flow cytometric analyses ashowing detection of endogenous PD-L1 expression on human 5637 urothelial cancer cells, either untreated (WT) or stimulated with IFN-γ (WT + IFN-γ), using AF647-conjugated PD-L1 quattrobinders compared to an AF647-conjugated anti-PD-L1 antibody. 5637 PD-L1 knockout (KO) cells are shown as a reference for specificity of the signals. **d** Quantification of the experiment described in b performed in biological triplicates (n=3). **e** Quantification of the experiment described in c performed in biological triplicates (n=3). **p<0.01; ***p<0.001; ****p<0.0001; two-way ANOVA with Šídák’s multiple comparisons test.

We then evaluated the ability of the PD-L1 quattrobinders to detect endogenous PD-L1 in the human urothelial carcinoma cell line 5637, which displays low basal PD-L1 expression that is upregulated upon IFN-γ stimulation. To confirm target specificity and rule out residual background, we generated a CRISPR-Cas9 engineered PD-L1 knockout cell line (5637 PD-L1 KO) and directly compared PD-L1 quattrobinder staining in wild-type, IFN-γ stimulated, and knockout cells (**Fig. 7c**). The PD-L1 quattrobinders exhibited no detectable background staining on 5637 PD-L1 KO cells. Wild-type 5637 cells exhibited robust labeling that was markedly enhanced after IFN-γ stimulation. Overall, intensity and staining patterns of the PD-L1 quattrobinders closely mirrored those of the anti-PD-L1 antibody in a robust and reproducible manner (**Fig. 7d,e**). Similar results were seen on the thyroid cancer cell line FTC-133, but BN_1339 showed background staining on the PD-L1 knockout cell line that was not observed for 5637 cells (Supplementary Fig. 8a). Unexpectedly, the BN_1358 quattrobinder produced weaker signals than other high-performing variants, despite its strong binding affinity (K_D_ = 2 nM) (**Fig. 3j**).

Given the inferior performance of our CD276 AI-binders relative to the PD-L1 binders across all assays tested so far, we also evaluated flow cytometry staining with their corresponding quattrobinders. The CD276 quattrobinders strongly labeled CHO cells overexpressing both the 2Ig and 4Ig CD276 isoforms, however, only BN_486-1519 produced a marginal signal in 5637 cells, and no staining was observed in SK-MEL-28 wild-type cells (Supplementary Fig. S8b–e). When benchmarked against a commercial anti-CD276 antibody, the signals generated by our CD276 quattrobinders were negligible. Furthermore, all our VTCN1 quattrobinders exhibited poor performance, with substantial background staining even on CHO cells (Supplementary Fig. 8f,g). Collectively, our data demonstrate that AI-quattrobinders can be used equivalent to conventional antibodies in flow cytometry and enable specific detection of endogenous protein expression levels, but their quality and specificity require rigorous validation and the efficiency to generate high quality AI-binders is dependent on the target.

## Discussion

The recent advances in AI-based protein design open new frontiers for biomedical research, including the generation of high-precision research tools and next-generation therapeutics. However, applicability and accessibility to a broader research community outside of groups specialized on protein design remains uncertain. Here, we show that combining our expertise in structural biology^32^, functional genomics^33^, and tumor immunology^34,35^ enables seamless incorporation of AI-based protein design into existing laboratory workflows, and leveraging it for the development of novel research tools adapted to our experimental needs. With regard to computational resources, the pilot pool was designed on a workstation with a single mid-range GPU (Nvidia RTX A5000) before employing a multi-GPU server with 8 RTX A5000 GPUs for the improved pools. Thus, our experience confirms that AI-based protein design can be leveraged even without the use of high-performance computation. However, resources become critical when scaling up or addressing more complex design tasks.

Pooled genetic screening in mammalian cells and phage display are established methods in many labs including ours^33,36,37^, which is why we integrated the RFdiffusion design pipeline with these screening techniques. The choice between mammalian cell-surface display and phage display should be guided by available expertise, research objectives, and intended applications of the AI-designed binders. The number of designs per target was informed by previously reported experimental success rates. Although DNA-oligo pool cloning can, in principle, accommodate very large libraries, we deliberately focused on libraries limited to 1,000–2,000 candidates to ensure a robust cell population for cell sorting. Future improvements to RFdiffusion and other protein design tools are expected to boost experimental success rates, enabling larger libraries and the multiplexing of more targets per screen. In this context, two recently released pipelines, BindCraft^8^ and AlphaProteo^7^, report high experimental validation rates for their de novo designed protein binders. BindCraft is an open-source tool which will facilitate rapid community-based confirmation. BindCraft designs are readily compatible with our screening platform, and a head-to-head comparison with RFdiffusion warrants future investigation. However, our results imply that it is beneficial to create a pool of AI-binders which can then be screened for the desired properties in subsequent experiments. Mere binding to the target protein is clearly important but, in many cases, not sufficient to create a usable research tool. Parameters such as binding kinetics, specificity, and stability are just as important but still difficult to reliably predict computationally.

Since the release of AlphaFold2^2^ and RoseTTAFold^4^, multiple structure prediction models are now openly available, including Chai-1^25^ and Boltz-1^26^. In our PD-L1 and CD276 binder datasets, the ipTM score generated by Chai-1 correlated well with experimental success and accurately predicted the majority of deleterious effects caused by binding interface mutations. Chai-1 integrates ESM language model embeddings with structural features to predict protein–protein interactions^25,28^. Unlike models reliant on multiple sequence alignments, the Chai-1 model captures sequence and structural context directly from single sequences, making it faster and well suited for predicting experimental success of AI-designed proteins that obviously lack evolutionary homologs. Nevertheless, the size of datasets is still limited, and accordingly our analysis is not an exhaustive comparison of the different algorithms. Furthermore, we used default parameters, and further parameter optimization could enhance the performance of the algorithms. As the number of datasets containing experimentally characterized AI-designed protein binders continue to grow, they will provide a valuable resource for benchmarking design algorithms and optimizing parameters. In addition, these datasets can be leveraged to fine-tune predictive models, specifically for AI-designed protein binders improving the accuracy of experimental success predictions.

Poor performing commercial antibodies remain an issue in biomedical research despite ongoing efforts to systematically evaluate their specificity and selectivity^29–31^. AI-designed protein binders offer a promising alternative, as their openly accessible sequences and structural models enable community-based validation and optimization. In this context, our study provides a proof of concept by demonstrating that PD-L1 quattrobinders can reliably detect endogenous protein levels on cancer cells by flow cytometry without appreciable background signals in most cases. Of course, we cannot exclude the possibility that the PD-L1 quattrobinders might exhibit further cross-reactivity and thus generate background signals in other cell lines not tested in our study, as seen in the case of BN_1339. However, we believe our findings outline a feasible strategy for developing reliable, open-source reagents for protein detection in flow cytometry and related applications. High yield and stability of AI-designed protein binders represent an additional advantage for their usage as research tools.

We see our study as an honest “field report” of non-experts in protein-design applying the available design algorithms to proteins of our interest, rather than proteins that produce the best success rates. We are convinced that the challenges encountered by us will be valuable for a broader research community aiming to implement AI-based protein design in their own projects. First, our CD276 AI-binders lack sufficient sensitivity to detect endogenous CD276 protein levels on cancer cells, both when incorporated into CAR-T cells and as quattrobinders. This limitation is likely due to weak binding affinity, considering that the previously published CD276 binder HM9 has picomolar affinity and successfully detects endogenous CD276 when deployed in CAR-T cells^21^. However, HM9 exhibits poor surface expression, at least in our hands, presumably because in vitro affinity maturation did not select for biochemical properties optimal for mammalian cell-surface expression. In contrast, our mammalian cell-surface display approach inherently selects AI-binders with robust expression and efficient trafficking to the cell surface.

Another limitation of our study is the poor specificity observed for our AI-binders targeting VTCN1, resulting in nonspecific background signals in flow cytometry. Clearly, specificity and selectivity of AI-binders remains a central challenge for their widespread use as antibody-like research reagents, and ultimately as therapeutics. We anticipate that future advancements in structure prediction^3,38^ and protein design algorithms^6–8^ will also enhance binder specificity and computational strategies will help to minimize or prevent cross-reactivity directly at the design stage.

In summary, we present a scalable workflow that bridges AI-based protein design with experimental validation across multiple prespecified targets, highlighting both successes and remaining challenges. Our results demonstrate that AI-designed binders can be functionally deployed in CAR-T cells and as quattrobinders for antibody-like flow cytometry applications. Together, these advances lay the groundwork for community-driven development of reliable, open-source protein detection tools for various biomedical applications, including flow cytometry. Our work also outlines the opportunities and challenges involved in translating AI-designed binders into biologics for cancer treatment.

## Methods

### AI-binder design pipeline

Our AI-binder design pipeline was implemented as a series of chained command-line steps (bash script). For each design job, an output directory was created and the input PDB, configuration file, and user-specified parameters (number of designs, contig sequences, hotspot residues) were set. RFdiffusion was then used with default parameters according to the instructions (https://github.com/RosettaCommons/RFdiffusion) to generate *de novo* poly-Gly backbone models. The DSSP program from the CCP4 suite^15,16^ was used to filter the designs and only structures with three or more helices were funneled into the next steps of the pipeline. PMPNN was used to design two alternative interface sequences for each poly-Gly backbone (https://github.com/nrbennet/dl_binder_design). Each design was scored with AlphaFold2 initial guess and only designs with a pAE interaction score < 10 were used in downstream experiments.

### Oligonucleotide pool design and cloning

Using our online pool design tool (jsb-lab.bio/AI-binders), AI-binder sequences were reverse-translated with random codon selection, omitting rare human codons and excluding randomly occurring Esp3I recognition sites. Overhangs, Esp3I sites, and common primer binding sites were added, and oligonucleotide sequences were filled up to the maximum oligonucleotide length using random DNA bases. Oligo sequences are provided in Supplementary Tables 2 and 5.

### DNA sequences

Primer sequences are provided in Supplementary Table 8.

### Phage display

To generate a library for phage display, the AI-binder library was amplified by PCR to introduce NotI and AscI restriction sites (primers fwd_NotI, rev_AscI) and the AI-binder library was cloned into a phagemid vector. The phage display was done using *E. coli* TG1 cells in combination with VCSM13 helper phages to enrich specific AI-binders. After phage production and purification by PEG/NaCl precipitation, the phages were precleared on magnetic streptavidin beads (Thermo Fisher 6560). The bait proteins were biotinylated by ChromaLink Biotin (Solulink B-1007-110) and immobilized to streptavidin beads. Two rounds of panning were performed using 20 µg and 2 µg of target protein for round one and two respectively. *E. coli* ER2738 cells were infected with the enriched phages and plated on LB-agar with 2YT/2 % glucose/Amp/Tet. Glycerol stocks were prepared of all libraries and plasmid mini-purification was performed. Plasmid DNA of phages display input, as well as output from two panning rounds were used as template for NGS library generation (primers NGS_phages_fwd and NGS_phages_rev, followed by a second PCR using Illumina barcode primers) and sequencing using the Illumina MiSeq platform.

### Library cloning for mammalian cell-surface display screening

The oligo pool (Twist Biosciences) was PCR amplified (primers pool-amplification_fwd, pool-amplification_rev) and cloned into the lentiviral target vector using Golden Gate assembly with the restriction enzyme Esp3I. The assembly was purified using a DNA Clean & Concentrator-25 column (Zymo) and was electroporated into 25 µL Endura electrocompetent *E. coli* (Biosearch Technologies) in 0.1-cm Bio-Rad cuvettes at 1,800 V, 10 µF, and 600 Ω. Cells were resuspended into 975 µL recovery medium and incubated for 1 hour at 37°C, 300 rpm. The suspension was then transferred into 450 mL LB medium containing 100 µg/mL ampicillin and grown overnight at 37°C, 180 rpm. Plasmid DNA was purified using the PureLink HiPure Plasmid Midiprep Kit (Invitrogen). AI-binder coverage of the plasmid library was determined by PCR amplification (primers NGS_cell-display_fwd and NGS_cell-display_rev, followed by a second PCR using DandJ barcode primers) and sequencing using the Illumina MiSeq platform.

### Library transduction for mammalian cell-surface display screening

With HBS and CaCl_2_, 1 × 10^6^ HEK293T cells were transiently transfected with the AI-binder plasmid library and the lentiviral packaging plasmids pMD2.G (Addgene, 12259) and psPAX2 (Addgene, 12260) at 2 µg, 0.65 µg, and 0.35 µg, respectively. The medium was changed after 16 and 24 hours. 40 hours post-transfection, the medium was collected and filtered through a 0.45 µm syringe filter. 2 × 10^6^ HEK293T cells were transduced with the filtered, virus containing supernatant at a 1:10 dilution aiming at transduction rates < 60 %. Transduced cells were selected by supplementing the medium with 1 µg/mL puromycin.

The HEK293T AI-binder library was harvested and incubated with respective Fc-tagged recombinant target proteins at 10 µg/mL (Supplementary Table 9) and secondary APC-labeled anti-human IgG Fc antibody at 250 ng/mL (clone M1310G05, Biolegend). Cells were sorted immediately after staining for distinct APC-signal using the fluorescence-activated cell sorter Sony MA900 at the Flow Cytometry Core Facility (FCCF), University of Bonn. Sorted cells were lysed in direct lysis buffer (1 mM CaCl2, 3 mM MgCl2, 1 mM EDTA, 1 % Triton X-100, 10 mM Tris pH 7.5) with proteinase K at 65 °C for 10 minutes and 95 °C for 15 minutes. Genomically integrated AI-binder sequences were PCR amplified using NEBNext High-Fidelity MasterMix (NEB) using primers pL_minibinder_fwd and pL_minibinder_rev. After second barcoding PCR (DandJ barcode primers), product purification, and NanoDrop-based quantification, AI-binders were sequenced on Illumina MiSeq or NextSeq platforms. Reads were filtered for a Q-Score > 30 and matched to AI-binder sequences. AI-binder counts were subjected to enrichment analysis (https://jsb-lab.bio/xyplot/).

### Individual AI-binder cloning and cell surface display validation

AI-binder sequences were ordered as individual gene fragments (Twist Biosciences) and cloned into the lentiviral target vector using Golden Gate assembly with the restriction enzyme Esp3I. Plasmid sequences were verified by Sanger sequencing (Microsynth).

HEK293T cells were transiently transfected with the AI-binder expression constructs using HBS and CaCl_2_. 36 to 48 hours post-transfection, cells were harvested and incubated with respective Fc-tagged target proteins at 10 µg/mL (Supplementary Table 9) and secondary APC-labeled anti-human IgG Fc antibody at 250 ng/mL, or with APC-labeled anti-HA.11 Epitope Tag antibody (clone 16B12, Biolegend). Cells were acquired at the analyzers BD FACSCanto II Flow Cytometer (FCCF Bonn) or 4-laser Cytek Aurora Spectral Flow Cytometer, and analyzed using FlowJo. For statistical analysis, the geometric mean of the APC-signal MFI of protein binding was normalized to the respective HA.11 epitope tag APC-signal to account for differences in surface presentation.

### Cell culture

The human cell lines HEK293T (fetal kidney, CVCL_0063), FTC-133 (thyroid carcinoma, CVCL-1219, kindly provided by Markus Essler, Nuclear Medicine, UKB), U87MG (glioblastoma, CVCL_0022), and their derivates were cultured in Dulbecco’s Modified Eagle Medium (DMEM) - GlutaMAX^TM^ medium (Gibco) supplemented with 10 % heat-inactivated FBS (Gibco) and 100 U/mL Penicillin-Streptomycin (Gibco). The human cell lines 5637 (bladder carcinoma, CVCL_0126), SK-MEL-28 (melanoma, CVCL_0526), SK-BR-3 (breast cancer, CVCL_0033), the Chinese hamster cell line CHO-K1 (ovary, CVCL_0214), and their derivates were cultured in Roswell Park Memorial Institute Medium (RPMI) 1640 - GlutaMAX^TM^ medium (Gibco) supplemented with 10 % heat-inactivated FBS (Gibco) and 100 U/mL Penicillin-Streptomycin. Platinum-E (Plat-E) cell lines were cultured in DMEM - GlutaMAX^TM^ medium, 10 % heat-inactivated FBS, 10 mM MEM Non-Essential Amino Acids Solution (Gibco), 100 U/mL Penicillin-Streptomycin, 1 mM Sodium Pyruvate (Gibco), and 10 mM HEPES (Roti). Primary human and murine cells were cultured in RPMI complete comprising RPMI 1640 - GlutaMAX^TM^ medium, 10 % heat-inactivated FBS, MEM Non-Essential Amino Acids Solution 1X (Gibco), 100 U/mL Penicillin-Streptomycin, 1 mM Sodium Pyruvate (Gibco), 10 mM HEPES (Roti), and 1X β- mercaptoethanol (Gibco), as well as below indicated additional experiment-specific supplements. All cell lines were cultured at 37 °C in 5 % CO_2_.

### Knockout (KO) cell line generation and validation

DNA templates for sgRNAs (Microsynth) against PD-L1, CD276, or VTCN1 were cloned into the CRISPR-Cas9 target vector px458 (Addgene, 48138) using Golden Gate assembly with the restriction enzyme BbsI. Plasmids were transfected into respective cell lines using FuGENE transfection reagent (Promega; 3:1 FuGENE reagent to DNA). 7 to 10 days post-transfection, cells were harvested and stained using fluorescently tagged antibodies against PD-L1 (clone IPI652.rMAb, BD Biosciences), CD276 (clone 185504, R&D Systems), or VTCN1 (clone MIH43, Biolegend), respectively. Cells were sorted for surface protein negative knock-out phenotype using the fluorescence-activated cell sorter Sony MA900 (FCCF, Bonn). Target sequences for sgRNA constructs were: PD-L1, CGCTGCATGATCAGCTATGG; CD276 AGGAAGATGCTGCGTCGGCG, and VTCN1, ATTGGAGACTTCCGAGAAGT.

### Generation of overexpression cell lines

Overexpression cell lines were created for the proteins PD-L1 (human Q9NZQ7-1, residues 1– 290), 2Ig CD276 (human Q5ZPR3-2, residues 1–316; Ig-like V-type domain 1 and Ig-like C-type domain 2), 4Ig CD276 (human Q5ZPR3-1, residues 1–534), and VTCN1 (human Q7Z7D3-1, residues 1–282). Gene fragments (Twist Biosciences) were cloned into the pRP233 or pRP236 target vectors (derived from Addgene plasmid #41841). HEK293T cells were transiently transfected with the protein expression construct and the retroviral packaging plasmids Gag-Pol and VSV-G at a ratio of 3:2:1. The medium was changed after 16 and 24 hours. 40 hours post-transfection, the virus-containing medium was collected, filtered through a 0.45 µm syringe filter, and immediately used for the transduction of CHO-K1, 5637, and SK-MEL-28 cells. Generated overexpression cell lines were sorted for purity using the fluorescence-activated cell sorter Sony MA900 (FCCF, Bonn). cDNA sequences cloned into pRP233 or pRP236 vectors are listed in Supplementary Table 8.

### Cloning of AI-binders for expression in *E. coli*

For expression of AI-binders in *E. coli*, coding sequences were subcloned from the respective lentiviral vector into the pEHISTEV vector^39^, which contains an N-terminal hexahistidine tag, followed by a tobacco etch virus (TEV) protease cleavage site. Cysteine point mutations were introduced into plasmids by primer-directed mutagenesis. For constructs with low expression yield, the coding sequences were codon-optimized for *E. coli* K-12 substr. MG1655 using the VectorBuilder Codon Optimization Tool (https://en.vectorbuilder.com/tool/codon-optimization.html) and synthesized as gene fragments (Twist Biosciences).

### Expression and Purification of AI-binders

Recombinant 6xHis-tagged AI-binders were expressed in *E. coli* BL21 (DE3) bacterial cells grown in LB medium containing 50 µg/mL kanamycin at 37 °C for 16 hours (pre-culture). The following day, larger culture volumes were inoculated with the pre-culture and adjusted to an optical density (OD_600_) of 0.1–0.2. Cultures were grown at 37 °C to an OD_600_ of 0.8 and protein expression was induced by addition of IPTG to a final concentration of 0.5 mM. Proteins were expressed at 18 °C for 16 hours. Bacteria were harvested by centrifugation at 4,000 × *g* for 25 minutes. Cell pellets were flash-frozen in liquid nitrogen and stored at -20 °C or subjected to immediate cell lysis.

Cell pellets were resuspended in lysis buffer (20 mM Tris pH 8.0, 500 mM NaCl, 20 mM imidazole) and lysed by sonication. Cell debris was sedimented by centrifugation at 25,000 × *g* for 45 minutes at 10 °C. The lysate was filtered through a membrane filter with a 0.8 µm pore size and subjected to affinity chromatography.

For affinity chromatography, Ni^2+^-NTA resin was equilibrated with lysis buffer and incubated with lysate for at least 2 hours at 4 °C or room temperature on a rolling incubator. The resin was transferred to a gravity column and washed extensively with lysis buffer and eluted with elution buffer (20 mM Tris pH 8.0, 500 mM NaCl, 500 mM imidazole). For removal of the 6xHis-tag, the elution fraction was dialyzed against dialysis buffer (20 mM Tris pH 8.0, 250 mM NaCl) for 16 hours at 4 °C and supplemented with 7xHis-tagged tobacco etch virus (TEV) protease (1:20 w/w) for 16 hours at 4 °C. For cysteine mutants, all buffers were additionally supplemented with 5 mM beta-mercaptoethanol.

For further purification following affinity chromatography or TEV digest, the AI-binders were subjected to size exclusion chromatography (SEC) on a HiLoad 16/600 Superdex 75 pg column (Cytiva) equilibrated in dialysis buffer. For SEC runs of cysteine mutants, the dialysis buffer was supplemented with 1 mM Tris(2-carboxyethyl)phosphine (TCEP). For TEV-digested proteins, the SEC column was equipped with an additional 5 mL HisTrap column (Cytiva) for removal of residual tag, non-cleaved protein and TEV protease.

### Biotinylation of AI-binders

AI-binders containing cysteine point mutations were biotinylated using the maleimide-reactive PEG_2_-Biotin label (EZ-Link, ThermoScientific). Prior to biotinylation, AI-binders were incubated in dialysis buffer containing 2 mM TCEP for 60 minutes at room temperature to reduce potential disulfide bonds. Immediately before addition of the biotin label, TCEP was removed by buffer exchange into labeling buffer (20 mM Tris or 20 mM HEPES pH 7.0, 250 mM NaCl) using 7K MWCO Zeba Spin desalting columns (ThermoScientific). The label was added at a 10-fold molar excess and incubated overnight at 4 °C. Excess label was removed by buffer exchange into labeling buffer using 7K MWCO Zeba Spin desalting columns.

### Quattrobinder assembly and flow cytometry staining

*E. coli* produced and biotinylated AI-binders were step-wise mixed and incubated with Alexa Fluor® 647–conjugated Streptavidin (Invitrogen) at a molar ratio of 4:1. Assembled quattrobinders were stored at -80 °C. For staining procedures, target cells were harvested and incubated with Zombie Violet™ Fixable Viability Kit (Biolegend) at 1:1,000 and quattrobinders at 150 nM. Samples were acquired at the analyzers BD FACSCanto II Flow Cytometer (FCCF Bonn) or 4-laser Cytek Aurora Spectral Flow Cytometer, and analyzed using FlowJo software (Tree Star).

### Surface plasmon resonance measurements

SPR measurements were performed on a Series S CM5 sensor chip (Cytiva) using a Biacore™ 8K instrument (GE Healthcare) operated at 25 °C. The chip was equilibrated in running buffer (20 mM HEPES pH 7.4, 150 mM NaCl, 1 mM TCEP, 0.05 % (*v/v*) Tween-20). Human PD-L1 extracellular domain (ECD) containing a C-terminal LPETG-tag and 6xHis-tag (hPD-L1) was immobilized by amine-coupling using 1:1 (*v/v*) mixture of 0.1 M N-hydroxysuccinimide (NHS) and 0.1 M 3-(*N*,*N*-dimethylamino)propyl-*N*-ethylcarbodimide (EDC) for surface activation. For immobilization, 0.5 µM hPD-L1 in 10 mM sodium acetate buffer (pH 4.5) was injected over the surface of the second flow cell for 600 seconds at a flow rate of 10 µL/min. Free binding sites on the surface were saturated with 1 M ethanolamine (pH 8) for 420 seconds at a flow rate of 10 µL/min. Single-cycle kinetics were measured with injections of increasing concentrations of the analyte over both flow cells. The association time was set to 120 seconds and the dissociation time to 600 seconds at a flow rate of 30 µL/min.

For the non-optimized and alanine-optimized variants of BN_PDL1_286, a five-fold concentration series ranging from 1.6 nM to 5000 nM was used. For BN_PDL1_482, a five-fold concentration series ranging from 0.32 nM to 1000 nM was used. For both variants of BN_PDL1_1358 and the alanine-optimized variant of BN_PDL1_482, a five-fold concentration series ranging from 0.064 nM to 200 nM was used.

Data were collected at a rate of 10 Hz, and double-referenced by reference flow cell subtraction and blank cycles. Binding parameters were obtained from the kinetic binding measurements using a 1:1 interaction model, using the Biacore^™^ Insight Evaluation Software (version 6.0.7.1750, Cytiva).

### Protein thermal stability analyses

Thermal stability of AI-binders was analyzed by nanoscale differential scanning fluorimetry using a Prometheus NT.48 (NanoTemper) device. Denaturation and renaturation of proteins was monitored by changes in internal fluorescence at wavelengths of 330 and 350 nm. Proteins were diluted to 10 µM in 20 mM HEPES pH 7.4, 150 mM NaCl. The changes in fluorescence were monitored from 20 °C to 90 °C (unfolding) followed by 90 °C to 20 °C (refolding) at a rate of 1.5 °C/min using the PR.ThermControl software (NanoTemper).

### Analytical size exclusion chromatography

Analytical size exclusion chromatography of AI-binders was performed on a Superose 6 3.2/300 column (Cytiva) equilibrated in SEC buffer (20 mM HEPES pH 7.4, 150 mM NaCl) and operated at a flow-rate of 0.05 mL/min using the Infinity II HPLC System (Agilent).

### Generation of AI-binder CAR constructs

The predicted AI binders were ordered as gene fragments (Twist Bioscience) and cloned into our second-generation CAR constructs using BbsI and Esp3I restriction sites. Our murine second-generation CAR comprised a CD8α leader sequence (Mouse NP_001074579.1, residues 1–27), AI binder, HA.11 epitope tag (YPYDVPDYA), CD8α hinge and transmembrane region (Mouse NP_001074579.1, residues 152-220), CD28z intracellular signaling domain (Mouse NP_031668.3, residues 178–218), and CD3z chain signaling domain (Mouse NP_001106862.1, residues 62–164). Similarly, the human second-generation CAR consisted of a CD8α leader sequence (Human NP_001139345.1, residues 1 - 21), AI binder, HA.11 epitope tag (YPYDVPDYA), CD8α hinge and transmembrane region (Human NP_001139345, residues 128 - 210), CD28z intracellular signaling domain (Human NP_006130.1, residues 180 - 220), and CD3z chain signaling domain (Human NP_000725.1, residues 52 - 163). Both human and murine 28z CAR were generated via gene synthesis and cloned into a pMSCV retroviral expression plasmid. CAR expression was coupled with a RQR8 reporter gene via a P2A sequence, as previously described^22^.

### Retroviral vector Production

Gamma retroviral supernatant for the transduction of primary human T cells was prepared as follows. HEK293T cells were transiently transfected with a mastermix containing Opti-MEM (Gibco), the respective AI-binder CAR plasmid (5 µg), packaging plasmids Gag-Pol (5 µg), RDF (3.33 µg), and FuGENE transfection reagent (3:1 FuGENE reagent to DNA) and cultured for 48 hours at 37 °C in 5 % CO_2_. Viral supernatant was filtered through 0.45 µM syringe filter and immediately used for the transduction of human T cells. For the generation of ecotropic gamma-retroviral supernatant for the transduction of primary murine T cells, Platinum-E (Plat-E) retroviral packaging cells were transiently transfected with a mastermix containing OptiMEM medium (Gibco), the respective AI-binder CAR plasmid (10 µg), and FuGENE transfection reagent (3:1 FuGENE reagent to DNA). Transfected cells were cultured at 37 °C in 5 % CO_2_. 48 hours post transfection the viral supernatant was harvested, filtered through 0.45 µM syringe filter and immediately used for the transduction of murine T cells.

### Human PBMCs

Peripheral blood from healthy donors was provided by the Institute for Experimental Hematology and Transfusion Medicine at the University Hospital Bonn, Bonn, Germany. All study procedures were performed in compliance with relevant laws and institutional guidelines and have been approved by the competent Ethics Committee of the Medical Faculty of the University of Bonn. Written informed consent was received prior to participation from the blood donors.

### Generation of CAR-T cells

To generate human CAR-T cells, peripheral blood mononuclear cells (PBMC) were thawed, resuspended in RPMI complete + 100 U/mL rhIL-2 (Peprotech) and activated for 48 hours at 37 °C and 5 % CO_2_ on CD3 and CD28 coated plates (1 µg/mL, OKT3, Biolegend; 3 µg/mL CD28.2, Biolegend). Next, 24-well non-tissue culture treated plates (Avantor) were pre-coated with 0.5 mL of 10 µg/mL RetroNectin (Takara Bio) at 4 °C overnight. Harvested retroviral supernatant was spun onto RetroNectin coated wells (at 2,000 x *g* for 2 hours at 32 °C). Subsequently, activated PBMCs in RMPI complete with 100 U/mL rhIL-2 were added and spinoculated at 800 x *g* for 90 minutes at 32 °C onto virus coated plates. T cells were subsequently cultured and expanded at a minimum density of 1 × 10^6^ in RPMI complete, supplemented with fresh rhIL-2 100 U/mL every 2 days. Cells were expanded for 7–10 days prior to use in functional assays. To generate murine CAR-T cells, splenocytes were harvested from C57Bl/6 mice, mechanically dissociated through a 70-μm filter and red blood cell (RBC) depleted using RBC lysis buffer (Hybri-Max, Sigma-Aldrich). Splenocytes were then resuspended in RMPI complete with 100 U/mL rhIL-2 (PeproTech) + anti-mCD28 (2 µg/mL, 37.51, Biolegend) and seeded onto CD3 (5 µg/mL 17A2, Biolegend) coated plates for 24 hours at 37 °C and 5 % CO_2._ Following activation, splenocytes were harvested and transduced via spinoculation (800 x *g* for 90 minutes at 32 °C) onto RetroNectin 10 µg/mL coated non-tissue culture treated plates covered with retroviral supernatant (2,000 x *g* for 2 hours at 32 °C). Splenocytes were subsequently expanded at a minimum density of 1 × 10^6^ in RPMI complete, supplemented with fresh 100 U/mL rhIL-2 for the first 48 hours. Afterwards, rhIL-2 was replaced with rmIL-7 (10 ng/mL) (PeproTech) and rmIL-15 (10 ng/mL) (PeproTech). Cells were expanded for 7–10 days prior to use in functional assays.

### Assessment of CAR-T cell transduction efficacy

CAR surface expression on primary T cells was analyzed via flow cytometry. Transduced cells were washed with 4 °C PBS, then blocked with Fc receptor blocking solution (TruStain FcX^TM^, Biolegend) and Live/Dead^TM^ Fixable Near-IR Dead Cell Stain Kit (Thermo Fischer) according to the manufacturer’s instructions. Cells were subsequently stained with fluorophore-coupled anti-hCD34 and anti-HA.11 antibodies to detect RQR8 and CAR expression on the surface of T cells, respectively. For the full surface staining, fluorophore-coupled antibodies against the following antigens were resuspended in FACS buffer (PBS, 2 % FBS, 2 mM EDTA): CD45, CD3, CD8, and CD4. Samples were acquired using the 4-Laser Cytek Aurora Spectral Flow Cytometer or 5-Laser Sony ID7000 Spectral Cell Analyzer, and analyzed using FlowJo software.

### CAR-T cell activation assay

CAR T cells were plated with indicated target cells at an effector to target ratio of 1:1 in 96-well U bottom plates in RPMI complete media for 5 hours, to determine intracellular cytokine production, or for 24 hours to measure the expression of surface CD25, at 37 °C and 5 % CO_2_. To assess IFN-γ production the protein transport inhibitors Monensin (GolgiStop^TM^, Becton, Dickinson and Company) and Brefeldin (GolgiPlug^TM^, Becton, Dickinson and Company) were added at 1:1,000. First, we performed a surface staining to identify CAR T cells as described above, followed by fixation and permeabilization for intracellular staining of IFN-γ and TNF-α according to the eBioscience^TM^ Foxp3/Transcription Factor Stain Buffer Set (Thermo Fischer). To assess induction of CD25 on CAR T cells, cocultures were harvested and surface staining was performed as described above including an anti-CD25 antibody. Samples were acquired with a 4-LaserCytek Aurora Spectral Flow Cytometer and analyzed using FlowJo software.

### Recombinant Fc-fusion protein expression and purification

The plasmid pD649-HAsp-CD276-Fc(DAPA)-AviTag-6xHis^40^ (Addgene #156861) was used for expression of the extracellular domain of 4Ig CD276 human Ig-Fc fusion protein. This plasmid was used as template to generate the construct encoding 2Ig CD276 human Ig-Fc fusion protein. 2Ig and 4Ig CD276 Fc constructs were produced and purified as previously described^41^. Briefly, 30 µg eukaryotic expression vector for 2Ig and 4Ig CD276 Fc constructs were transiently transfected into HEK293F cells and recombinant 2Ig and 4Ig CD276 Fc-fusion proteins were purified under sterile conditions from culture supernatants using Protein G beads (GE Healthcare). Glycine solution (0.1 M, pH 3.0) was used for elution from Protein G beads, and the solution was buffered to pH 7.4 using Tris buffer (1 M, pH 8). Concentrations were determined by ELISA. pD649-HAsp-CD276-Fc(DAPA)-AviTag-6xHis was a gift from Chris Garcia (Addgene plasmid #156861; http://n2t.net/addgene:156861; RRID:Addgene_156861)^40^. Commercially available Fc-fusion proteins, or provided by the Core Facility Nanobodies (CFN, UKB, Bonn), used in this study are listed in Supplementary Table 9.

### Rescoring

To score the designed structures in complex with their target we used AlphaFold 2 (AF2) initial guess^13^, Chai-1^25^ and Boltz-1^26^. All models were run on a single Nvidia L40S GPU. AF2 initial guess was used through the scripts provided by Bennet et al.^13^ and run with default parameters on all designs. pAE interaction scores were calculated after the initial design and filtering over the number of alpha-helices. For further rescoring, we assessed the results that were included in the AI-binder design libraries. Chai-1 (v.0.5.2) was used via the Python interface provided by the authors^25^. The model was run with sequence information and ESM2 embeddings as input. Sequences were obtained from the PDB files that were outputted from the AF2 initial guess pipeline. No multiple sequence alignments (MSAs) were provided to the model. For scoring we decided on using the standard settings: 3 truck recycles, 200 diffusion time steps. Boltz-1 (v.0.4.0) was run with recycling_steps=10, diffusion_samples=5 as input parameters. MSAs were generated to provide evolutionary information for subsequent protein structure prediction. Protein sequences were compiled into a single FASTA format file. Each sequence contained a unique identifier in the header line followed by the amino acid sequence. Two complementary sequence databases were employed for comprehensive homology searching: uniref30_2302_db and colabfold_envdb_202108_db. MSAs were generated using the MMseqs2 suite with GPU acceleration^27^ through ColabFold^42^. The specific procedure was as follows: MMseqs2 GPU servers were initialized for both the UniRef30 and environmental databases with the following parameters:

- Maximum sequences per query: 10,000

- Database load mode: 0

- Prefilter mode: 1

Sequence similarity searches were performed using the ColabFold search tool with these key parameters:

- –gpu 1

- –gpu-server 1

- -s 8.5

- –db-load-mode 0

- –pairing_strategy 1

- –use-env 1

- –use-templates 0

For each query sequence in the input FASTA file, homologous sequences were identified, aligned, and compiled into separate MSA files in A3M format. The quality of each MSA was assessed by quantifying the number of sequences captured, providing an indication of the depth of evolutionary information available for each query protein. The entire MSA generation process was executed in an automated fashion using a custom pipeline implemented in Bash. These MSAs were subsequently utilized as input for protein structure prediction via Boltz-1.

All used models and packages are available on GitHub under the following repositories: AF2 initial guess (https://github.com/nrbennet/dl_binder_design), Chai-1 (https://github.com/chaidiscovery/chai-lab), Boltz-1 (https://github.com/jwohlwend/boltz), ColabFold (https://github.com/sokrypton/ColabFold). Databases for MSA generation were obtained from https://colabfold.mmseqs.com.

## Data availability

PDB files of designs will be available in Zenodo.

## Code availability

Code for the adjusted RFdiffusion pipeline and rescoring will be available on GitHub https://github.com/HoelzelLab/IEO_AI_Binder_cancer_surface_2025.

## Acknowledgements

We would like to thank the Flow Cytometry Core Facility (FCCF) and the Core Facility Nanobody (CFN) of the Medical Faculty at the University of Bonn for providing support and instrumentation funded by the Deutsche Forschungsgemeinschaft (DFG, German Research Foundation). We also thank Sally Liu, Fenna Floortje Feenstra, and Saskia Vadder (IEO, University Hospital Bonn) for their excellent technical assistance and help.

D.F. was supported by a fellowship from the Studienstiftung des deutschen Volkes, Bonn, Germany. S.K. is supported by the Bavarian Cancer Research Center (BZKF) (TANGO), the Deutsche Forschungsgemeinschaft (DFG, grant number: KO5055-2-1 and KO5055/3-1), the international doctoral program ‘i-Target: immunotargeting of cancer’ (funded by the Elite Network of Bavaria), the Melanoma Research Alliance (grant number 409510), Marie Sklodowska-Curie Training Network for Optimizing Adoptive T Cell Therapy of Cancer (funded by the Horizon 2020 programme of the European Union; grant 955575), Marie Sklodowska-Curie Training Network for tracking and controlling therapeutic immune cells in cancer (funded by the Horizon Programme of The EU, grant 101168810), Else Kröner-Fresenius-Stiftung (IOLIN), German Cancer Aid (AvantCAR.de), the Wilhelm-Sander-Stiftung, Ernst Jung Stiftung, Institutional Strategy LMUexcellent of LMU Munich (within the framework of the German Excellence Initiative), the Go-Bio-Initiative, the m4-Award of the Bavarian Ministry for Economical Affairs, Bundesministerium für Bildung und Forschung, the EUROSTAR-Programm, European Research Council (Starting Grant 756017, PoC Grant 101100460 and CoG 101124203), by the SFB-TRR 338/1 2021–452881907, Fritz-Bender Foundation, Deutsche José Carreras Leukämie Stiftung, Hector Foundation, Bavarian Research Foundation (BAYCELLATOR), the Monika-Kutzner Foundation, the Bruno and Helene Jöster Foundation (360° CAR), the Dr. Rurainski-Foundation. A.S. was supported by the Federal Ministry of Education and Research (BMBF) within the framework of the funding programme ACCENT (funding code 01EO2107). T.R. was funded by the Deutsche Forschungsgemeinschaft (DFG) (Emmy Noether Grant Project-ID 506620580). M.H. is a member of the CANTAR project, which receives funding from the Netzwerke 2021 program, an initiative of the Ministry of Culture and Science of the State of North Rhine-Westphalia. M.H. is supported by the Deutsche Krebshilfe (German Cancer Aid) project grant 70114292 (Excellence Program for established scientists) and within SFB 1399 by the Deutsche Forschungsgemeinschaft (DFG) – Project ID 413326622. J.L.S.-B. was supported by the Hans und Ria Messer Stiftung. M.G. is supported by the European Research Council (ERC Advanced Grant NalpACT). M.I.T., T.R., M.G., T.B., J.L.S.-B., G.H. and M.H. are members of ImmunoSensation – the immune sensory system – supported by the Deutsche Forschungsgemeinschaft under Germany’s Excellence Strategy EXC2151 - project ID 390873048. The sole responsibility for the content of this publication lies with the authors.

## Author contributions

Acquisition of data: B.B., S.C.B., B.A.ME, C.I.F., J.M.M., E.T., D.F., M.C.R.Y., M.K., A.H., K.B., J.M.P.T., S.B., K.M. Data analysis and visualization: B.B., S.C.B., B.A.ME, C.I.F., J.M.M., E.T., D.F., M.C.R.Y., A.H., S.M., T.B., J.L.S.B, G.H., M.H. Resources: M.I.T., A.S., S.K., J.O., H.R., T.R., M.G., S.M., T.B., J.L.S.B, G.H., M.H. Coding: T.N.K., G.H., M.H., J.L.S.B., B.B. Protein designs: G.H., M.H. Conceptualization: S.M., T.B., J.L.S.B, G.H., M.H. Reviewing and editing: B.B., S.C.B., B.A.ME, C.I.F., J.M.M., S.K., M.G., S.M., T.B., J.L.S.B, G.H., M.H. Writing: G.H., M.H. All authors read and approved the final manuscript.

## Supplementary Figure Legends

**Supplementary Figure 1.**
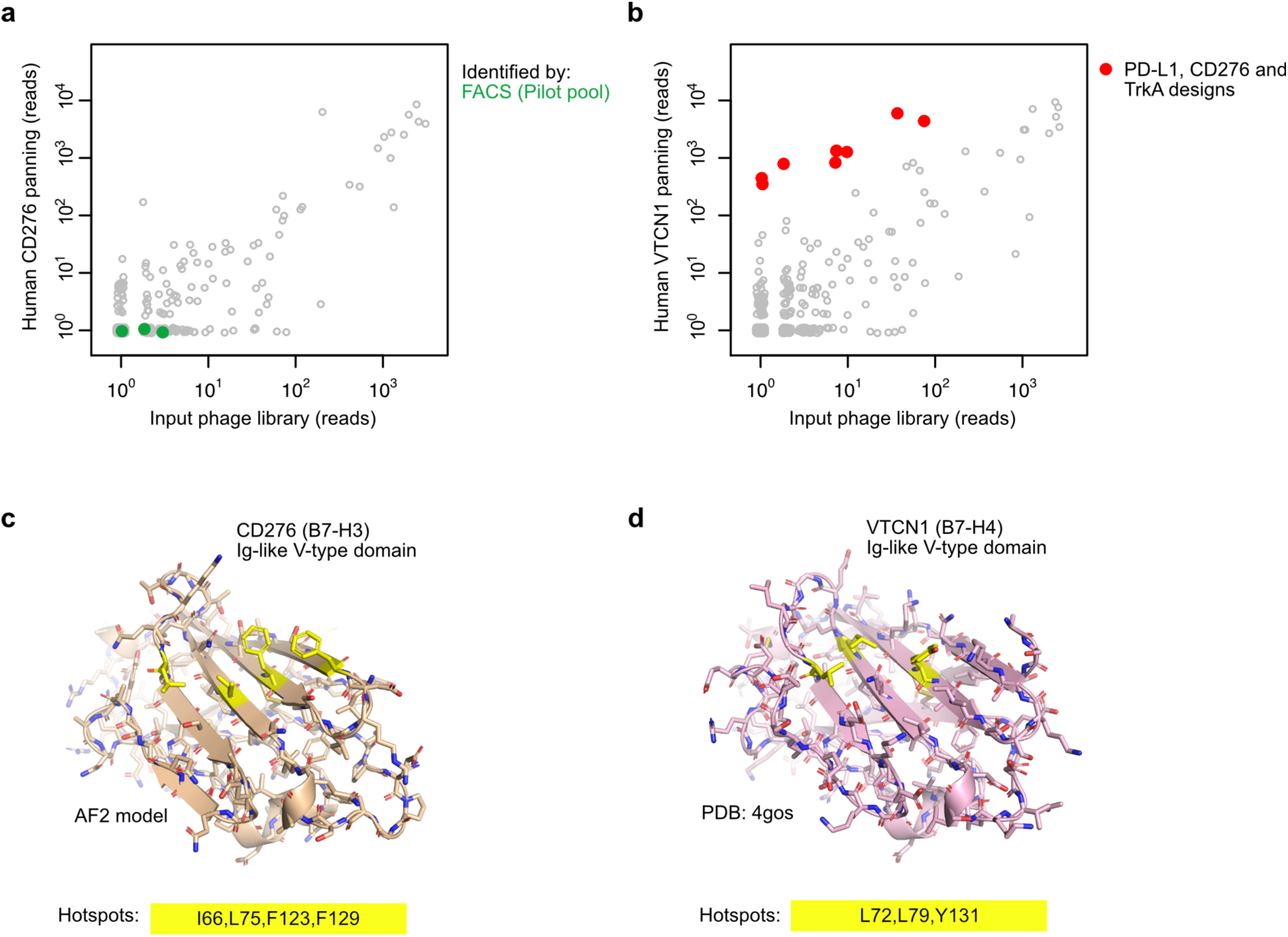
CD276 and VTCN1 phage display and hotspot selection for improved pool design. **a** Scatter plot summarizing the NGS results from phage display and panning with human CD276 protein. **b** As described in a but for panning with human VTCN1 protein. Modest enriched candidates were identified as off-target binders. **c** Cartoon representation with side chains of human CD276 structure (AF2 model). Selected hotspot residues for RFdiffusion design pipeline highlighted in yellow as used for design of the improved pool. **d** As described in c but for VTCN1 (PDB:4gos).

**Supplementary Figure 2.**
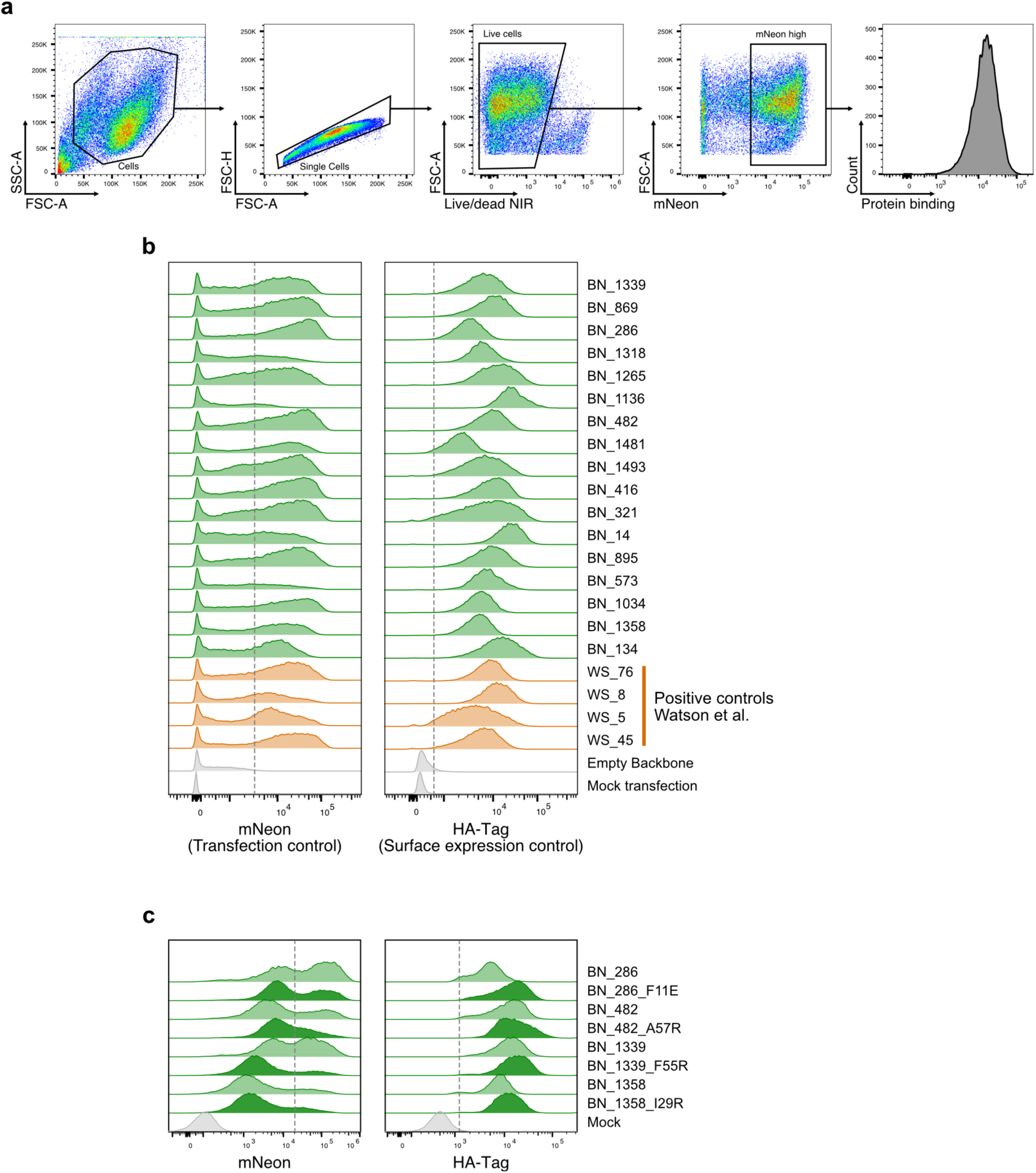
PD-L1 AI-binder validation: Transfection and surface expression controls. **a** Representative FACS plots showing gating strategy. **b** Flow cytometric analyses showing proportions of HEK293T cells expressing PD-L1 AI-binders by mNeon positivity and cell surface expression of AI-binders by HA-tag signal. **c** As described in b but for PD-L1 AI-binder interface mutants.

**Supplementary Figure 3.**
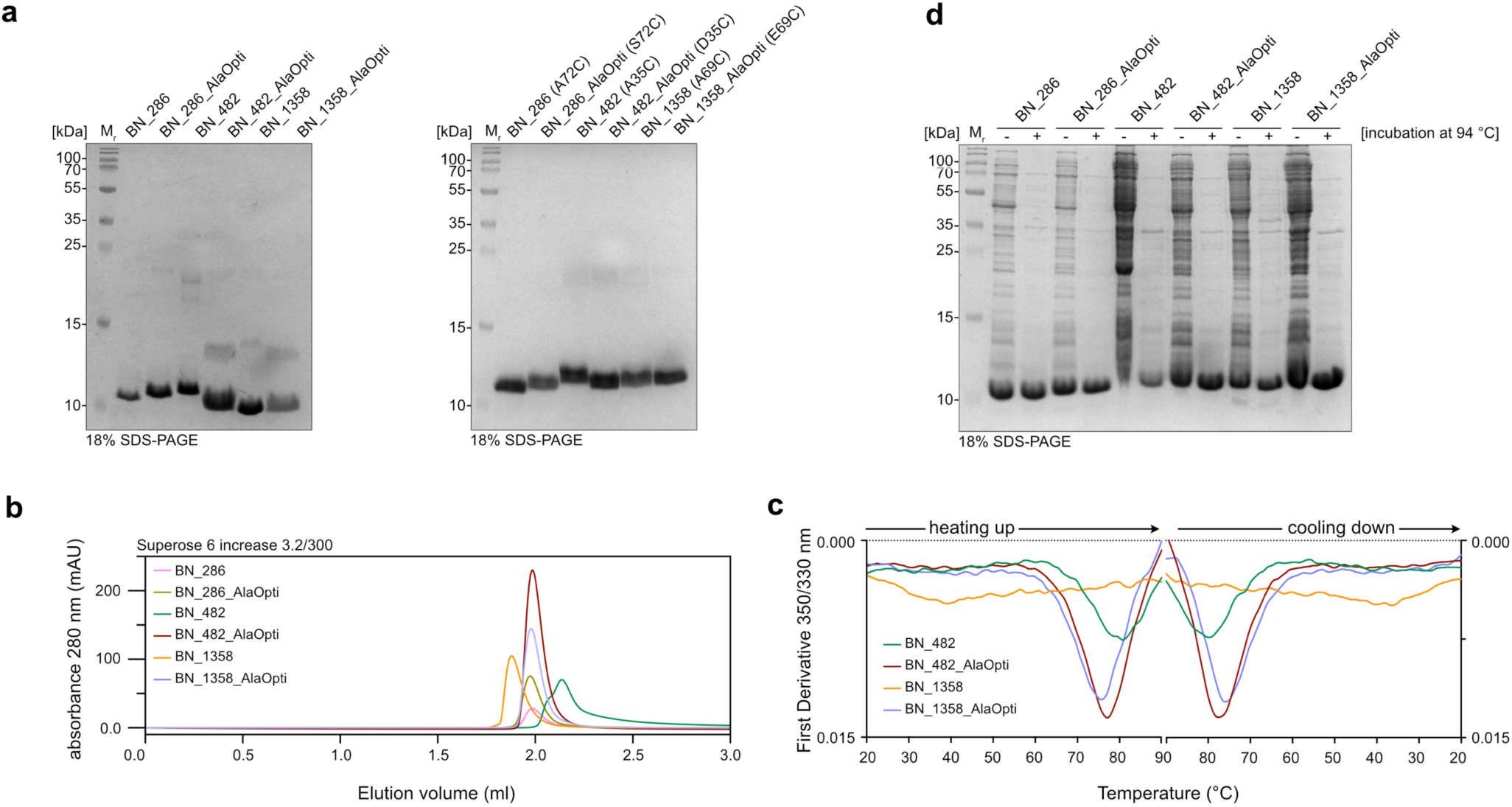
Biochemical analysis of AI-binder expression and analysis of the thermal stability. **a** Coomassie-stained SDS-PAGE of PD-L1 specific AI-binders (left panel) or PD-L1 specific AI-binders containing single-cysteine mutations (right panel) purified from *E. coli*. **b** Superpositions of analytical gel filtration runs of selected PD-L1 specific AI-binders using a Superose 6 3.2/300 column. **c** Nano differential scanning fluorometry (nanoDSF) experiments of selected PD-L1 specific AI-binders. The y-axis shows the first derivative of the corresponding melting curve. The two panels show the unfolding (left) and refolding (right) stages of the experiment. **d** Coomassie stained SDS-PAGE of *E. coli* lysates after the overexpression of AI-binders with (+) and without (-) incubation at 94 °C for 20 min. The boiled samples were centrifuged and the cleared supernatant was loaded onto the gel.

**Supplementary Figure 4.**
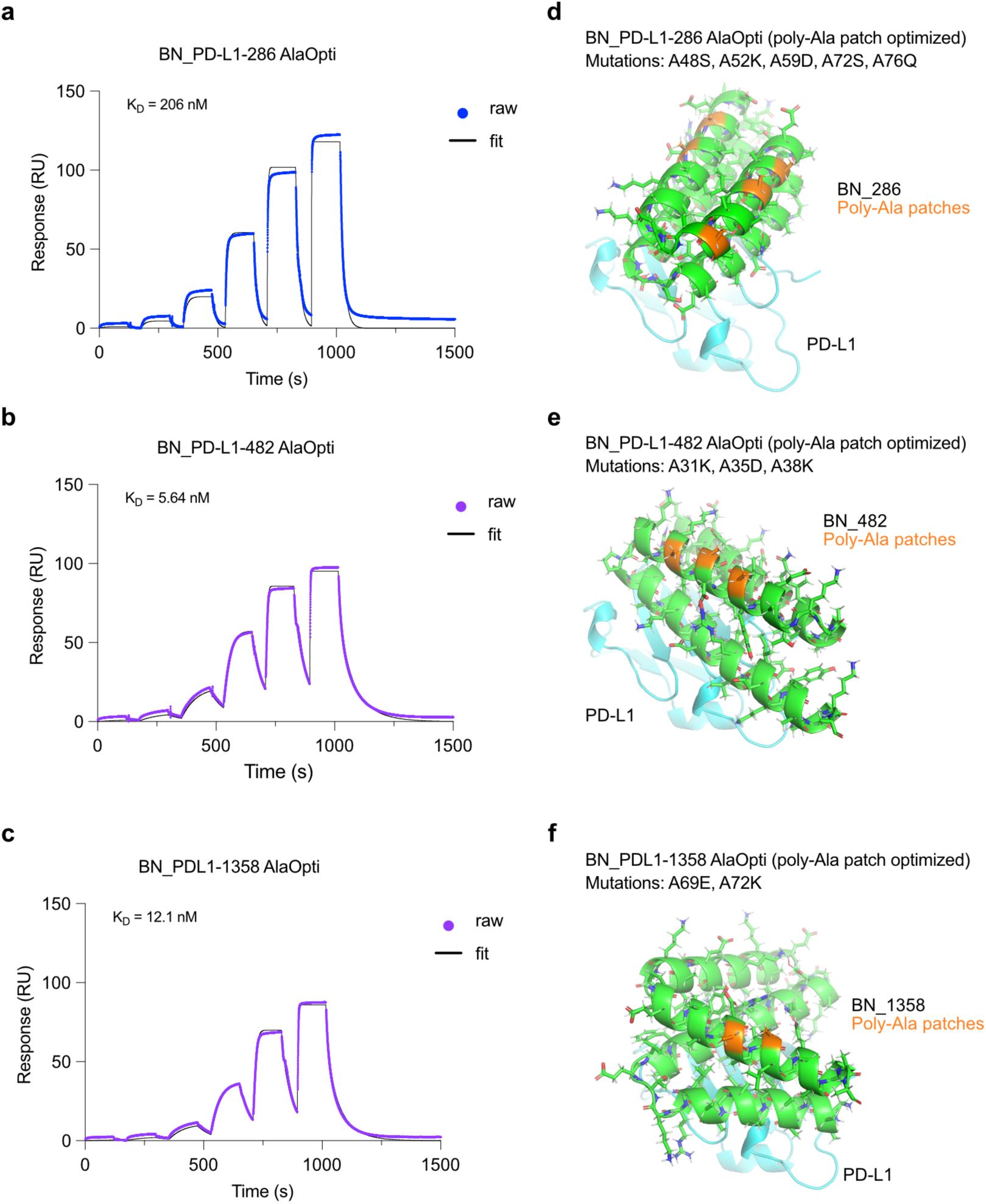
Binding affinity measurements of PD-L1 AI-binders with optimization of poly-alanine patches. **a** SPR (Surface Plasmon Resonance) sensorgrams showing raw binding data (colored dots) and fitted curves (black) used to determine binding affinities (K_D_) of poly-Ala patch optimized (AlaOpti) PD-L1 AI-binders BN_286, **b** BN_482, and **c** BN_ 1358. **d** Cartoon representation with side chains as predicted by AF2 model in RFdiffusion pipeline showing poly-Ala patches of BN_286 and the introduced mutations to generate AlaOpti variant. **e** Same as described in d but for BN_482 and **f** BN_1358.

**Supplementary Figure 5.**
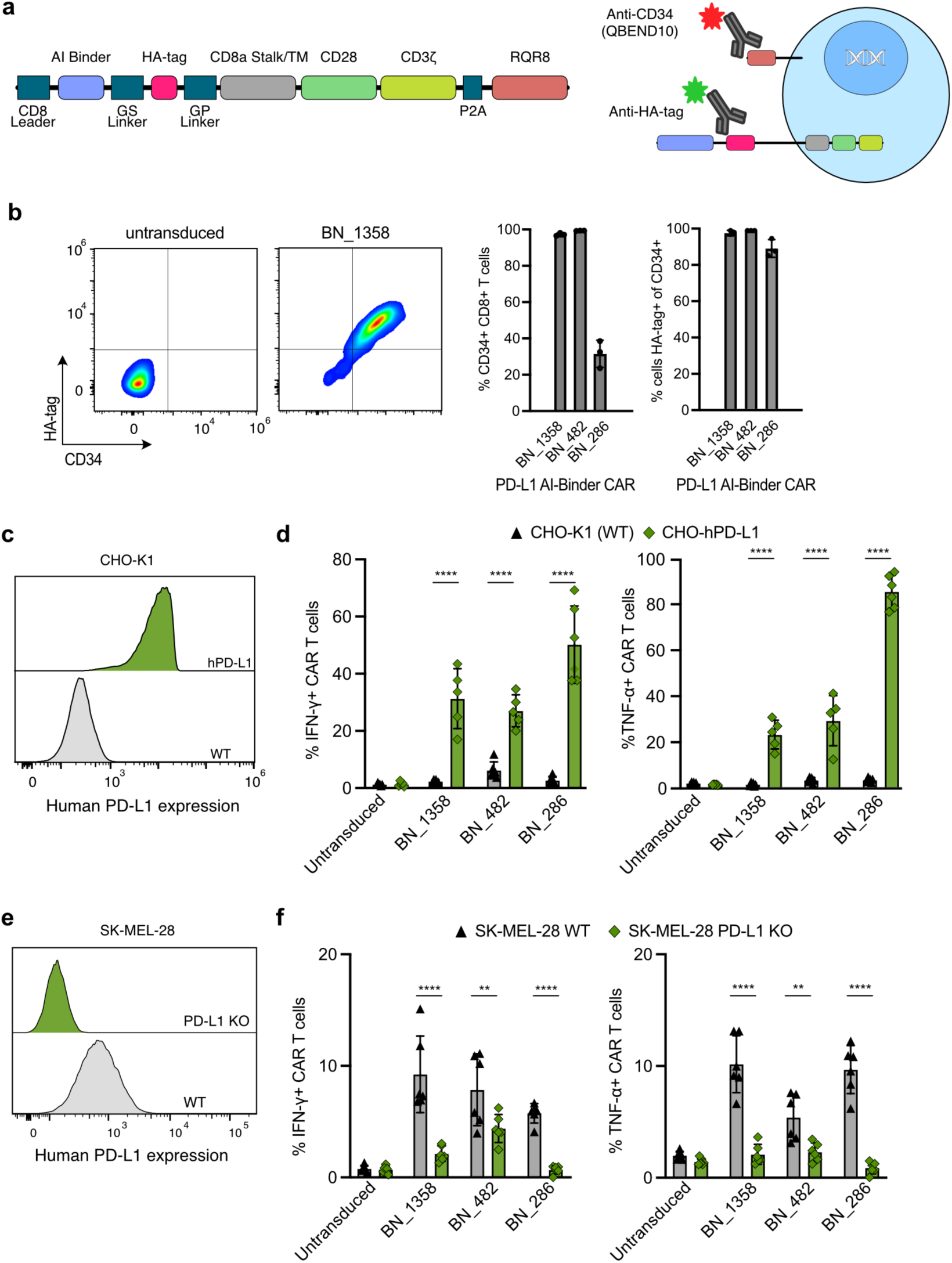
PD-L1 AI-binder CAR-T cells specifically recognize endogenous PD-L1 expressed on cancer cells. **a** Outline of deployed second-generation CAR construct and flow cytometric strategy to assess transduction efficiency (RQR8) and CAR cell surface expression (HA-tag). **b** Representative flow cytometric plots showing successful transduction and cell surface expression of PD-L1 AI-binder CAR (BN_1358) on T cells and corresponding quantification (n=3 biological replicates) with three PD-L1 AI-binders. **c** Representative flow cytometry histograms showing overexpression of human PD-L1 in CHO cells transduced with a construct encoding for human PD-L1 (hPD-L1) versus control CHO cells (CHO-K1). **d** Activation of PD-L1 AI-binder murine CAR-T cells exposed to CHO-hPD-L1 cells or CHO-K1 control cells determined by flow cytometric analyses of intracellular cytokine (IFN-γ, TNF-α) expression (n=3). **e** Representative flow cytometry histograms showing PD-L1 surface expression on human SK-MEL-28 wild-type (WT) cells and CRISPR-Cas9 engineered SK-MEL-28 PD-L1 knockout (KO) cells. **f** Activation of PD-L1 AI-binder murine CAR-T cells exposed to SK-MEL-28 WT cells and SK-MEL-28 PD-L1 KO cells determined by flow cytometric analyses of intracellular cytokine (IFN-γ, TNF-α) expression. Quantification of experiments performed in biological triplicates (n=3). **p<0.01; ***p<0.001; ****p<0.0001; two-way ANOVA with Šídák’s multiple comparisons test.

**Supplementary Figure 6.**
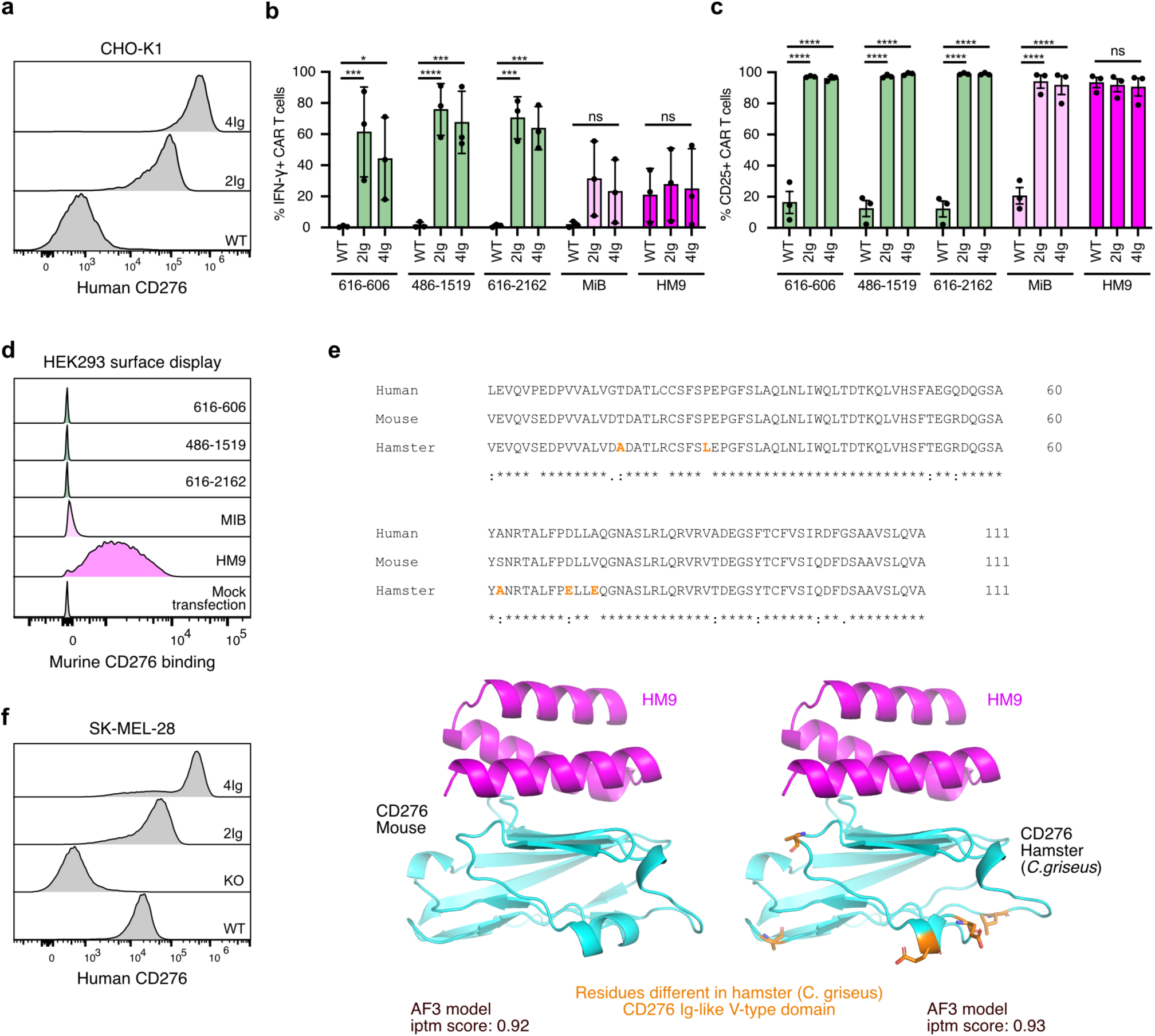
CD276 AI-binder CAR-T cells recognize overexpressed CD276 on CHO cells and show different profiles of species cross-reactivity. **a** Representative flow cytometry histograms showing surface expression of 2Ig and 4Ig CD276 isoforms overexpressed on CHO cells compared to untransduced controls (WT). **b** Activation of CD276 AI-binder human CAR-T cells exposed to CHO cells expressing 2Ig CD276, 4Ig CD276 or control cells determined by flow cytometric analyses of intracellular cytokine (IFN-γ) expression. Quantification of experiments performed in biological triplicates (n=3). **c** Same as described in b but for induced CD25 surface expression on activated human CAR-T cells (n=3). **d** Representative flow cytometry histograms showing murine CD276 binding to CD276 AI binders expressed on HEK293T cells. **e** Protein sequence alignment of the Ig-like V-type domain of CD276 from human, mouse, and hamster (*C. griseus*). Below, cartoon representations of AF3 models showing the HM9 CD276 AI-binder in complex with murine and hamster CD276. Residues differing between murine and hamster CD276 are highlighted in orange. AF3 ipTM scores are indicated. **f** Flow cytometry histograms showing endogenous CD276 surface expression on human SK-MEL-28 melanoma cells (WT), CRISPR-Cas9 engineered SK-MEL-28 CD276 knockout (KO) cells, and 2Ig and 4Ig CD276 isoforms overexpressed SK-MEL-28 KO cells on. *p<0.05; ***p<0.001; ****p<0.0001; two-way ANOVA with Šídák’s multiple comparisons test.

**Supplementary Figure 7.**
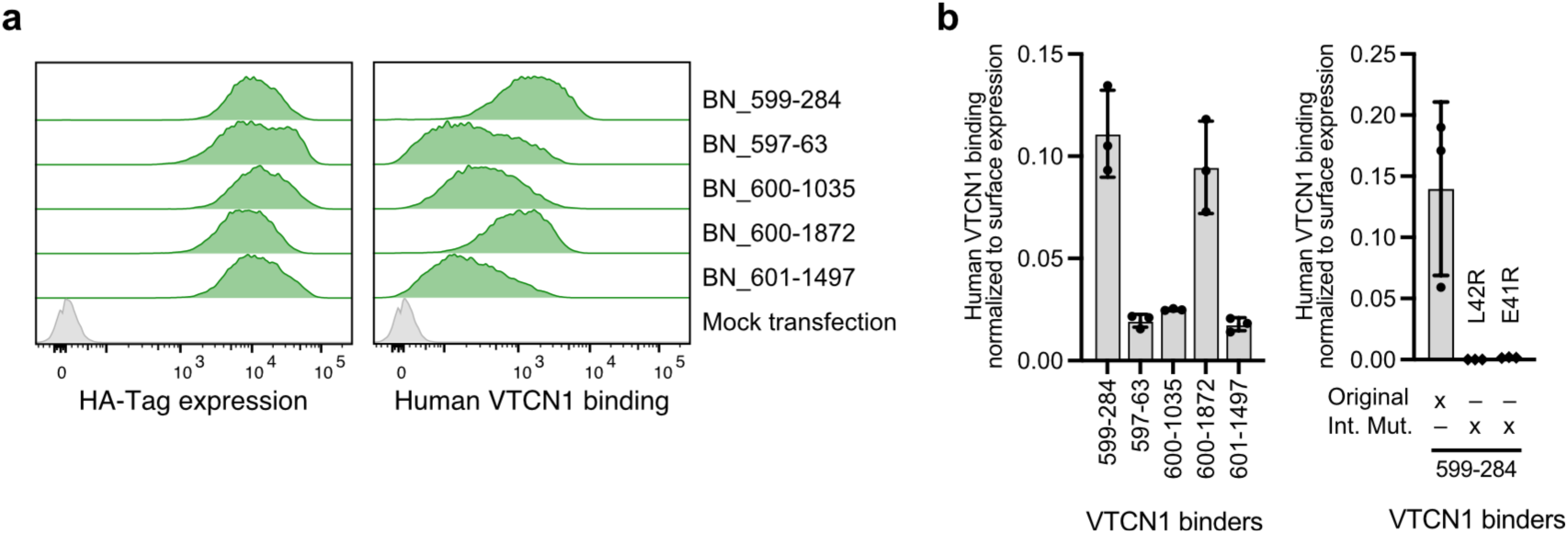
Validation of VTCN1 AI-binders. **a** Flow cytometry histograms showing cell surface expression of VTCN1 AI-binders on HEK293T cells and binding of human VTCN1 Fc-fusion protein. **b** Quantification of experiment described in a performed in biological triplicates (n=3), including the analyses of VTCN1 interface mutants.

**Supplementary Figure 8.**
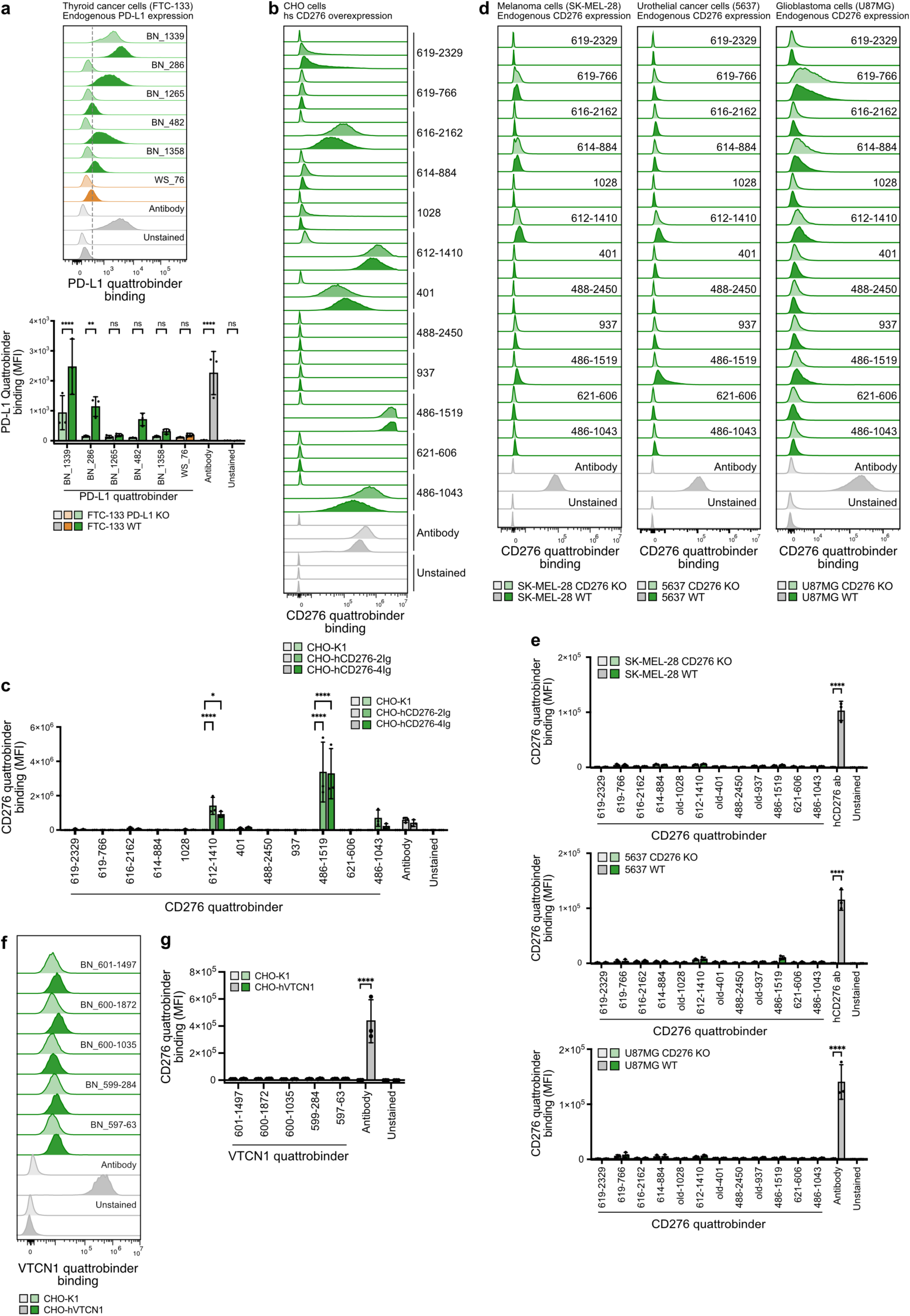
Evaluation of CD276 and VTCN1 quattrobinders. **a** Representative flow cytometry histograms showing staining with Alexa Fluor® 647 (AF647)-conjugated PD-L1 quattrobinders compared to an AF647-conjugated anti-PD-L1 antibody of thyroid cancer FTC-133 wild-type (WT) cells and CRISPR Cas9-engineered PD-L1 knockout (KO) cells. Quantification of flow cytometry data, performed in biological triplicates (n=3). **b** Representative flow cytometry histograms showing detection of human 2Ig and 4Ig CD276 overexpressed on CHO cells (CHO-hCD276-2Ig, -4Ig) compared to wild-type CHO (CHO-K1) cells by AF647-conjugated CD276 quattrobinders and an AF647-conjugated anti-CD276 antibody. **c** Quantification of experiment described in b, performed in biological triplicates (n=3). **d** Representative flow cytometry histograms showing staining with AF647-conjugated CD276 quattrobinders compared to an AF647-conjugated anti-CD276 antibody of CD276 wild-type (WT) and CRISPR Cas9-engineered CD276 knockout (KO) cancer cells (human SK-MEL-28 melanoma cells; human 5637 urothelial cancer cells; human U87MG glioblastoma cells). **e** Quantification of experiments described in d, performed in biological triplicates (n=3). **f** Representative flow cytometry histograms showing staining with AF647-conjugated VTCN1 quattrobinders compared to an APC-conjugated anti-VTCN1 antibody of human VTCN1 overexpressed on CHO (CHO-hVTCN1) cells and wild-type CHO (CHO-K1) cells. **g** Quantification of experiment described in f, performed in biological triplicates (n=3). **p<0.01; ****p<0.0001; two-way ANOVA with Šídák’s multiple comparisons test.

