## Supplementary_Table_9 for "Development of AI-designed protein binders for detection and targeting of cancer cell surface proteins"

**Supplementary Table 9. List of Fc-fusion proteins either commercially available or provided by the core facility nanobody (University Hospital Bonn).**

| **Protein** | **Amino acids and accession number** | **Vendor** | **Catalogue number** | **Used for** |
| --- | --- | --- | --- | --- |
| Human PD-L1  (llama Fc-tag) | xxx | Core Facility Nanobody | – | Screen of pilot pool |
| Human PD-L1  (human Fc-tag) | AA Phe 19 – Arg 238  Accession #NP_054862.1 | ACROBiosystems | PD1-H5258 | Screen of pilot pool  Screen of improved pool  Validation of both pools |
| Mouse PD-L1  (human Fc-tag) | AA Phe 19 – Thr 238  Accession #NP_068693 | ACROBiosystems | PD1-M5251 | Screens of both pools  Validation of both pools |
| Human PD-L1  (LPETG, 6xHis-tag) | AA Ala 18 – Arg 238  Accession #NP_054862.1 | Core Facility Nanobody | – | SPR spectroscopy |
| Human CD276 (2Ig)  (human Fc-tag) | AA Leu 29 – Pro 245  Accession#Q5ZPR3-2 | ACROBiosystems  Antibodies Online | B73-H5253  ABIN7273959 | Screen of pilot pool  Screen of improved pool  Validation of both pools |
| Human CD276 (4Ig)  (human Fc-tag) | AA Gly 27 – Thr 461  Accession # Q5ZPR3-1 | ACROBiosystems | B7B-H5258 | Validation of both pools |
| Mouse CD276  (llama Fc-tag) | xxx | Core Facility Nanobody | – | Screen of pilot pool |
| Mouse CD276  (human Fc-tag) | AA Val 29 – Phe 244  Accession #NP_598744 | ACROBiosystems | B73-M5255 | Validation of both pools |
| Human VTCN1  (llama Fc-tag) | xxx | Core Facility Nanobody | – | Screen of pilot pool |
| Human VTCN1  (human Fc-tag) | AA Phe 29 – Ala 258  Accession #Q7Z7D3-1 | ACROBiosystems | B74-H5256 | Screen and validation of improved pool |
